# Synergistic anti-phage activity of bacterial defence systems

**DOI:** 10.1101/2022.08.21.504612

**Authors:** Yi Wu, Sofya K. Garushyants, Anne van den Hurk, Cristian Aparicio-Maldonado, Simran Krishnakant Kushwaha, Claire M. King, Yaqing Ou, Thomas C. Todeschini, Martha R.J. Clokie, Andrew D. Millard, Yilmaz Emre Gençay, Eugene V. Koonin, Franklin L. Nobrega

## Abstract

Bacterial defence against phage predation involves diverse defence systems acting both individually and concurrently, yet their interactions remain poorly understood. We investigated >100 defence systems in 42,925 bacterial genomes and identified numerous instances of their non-random co-occurrence and mutual exclusion. For several pairs of defence systems significantly co-occurring in *Escherichia coli* strains, we demonstrate synergistic anti-phage activity, including tmn synergising with defence systems containing sensory switch ATPase domains. Some of the defence system pairs that are mutually exclusive in *E. coli* significantly co-occur in other taxa, and one tested mutually exclusive pair showed synergy. These findings demonstrate compatibility and synergy between defence systems, allowing bacteria to adopt flexible strategies for phage defence, driven by specific environmental pressures. The evolution of bacterial immune repertoires seems to be shaped largely by selection for resistance to host-specific phages rather than by negative epistasis.

## INTRODUCTION

Bacteria evolved numerous, diverse lines of active immunity as well as abortive infection mechanisms to withstand phage predation^1^. Recent systematic screening uncovered numerous anti-phage defence systems that widely differ in protein composition and modes of action^2–7^. The mechanisms employed by bacterial defence systems include phage genome or protein sensing followed by degradation^8–10^, introduction of modified nucleotides that abrogate phage replication^11,12^, as well as multiple sensing mechanisms leading to abortive infection that results in the host cell dormancy or death^4,13–21^. However, for many, perhaps, the majority of the bacterial defence systems, the mechanism of action remains unknown.

A bacterial genome carries, on average, about five distinct (currently identifiable) defence systems^22^. The remarkable variability of immune repertoires was observed even within the same species^22–24^. Genes encoding components of these systems tend to cluster together in specific genomic regions known as defence islands, sometimes associated with mobile genetic elements (MGEs) integrated into distinct hotspots in the bacterial genome^24–26^. Defence systems are often horizontally transferred between bacteria, and close proximity of the respective genes could facilitate simultaneous transfer of multiple systems.

Despite the recent burst of bacterial defence system discovery, the causes of their clustering in defence islands remain poorly understood. It has been argued that co-localisation of defence systems in MGEs and the resulting joint horizontal gene transfer (HGT) could provide fitness advantages to recipient bacteria, especially in phage-rich environments^27^. Additionally, it has been suggested that synergistic interactions between defence systems could drive their co-localization and favour their joint transfer^28,29^, as supported by the conservation of certain sets of defence systems^30^. For example, CRISPR-Cas systems of different subtypes often co-occur and the CRISPR arrays interact with Cas proteins across different systems^31^. Furthermore, toxin-antitoxin (TA) RNA pairs^32^ and possibly other TA modules^33^ safeguard CRISPR immunity by making cells dependent on CRISPR-Cas for survival. CRISPR-Cas and restriction-modification (RM) systems^34^, as well as BREX and the restriction enzyme BrxU^29^, co-occur resulting in expanded phage protection. However, these examples of interaction between bacterial defence systems notwithstanding, their co-occurrence in bacteria and the connections between co-occurrence and co-localization in bacterial genomes have not been analysed on a large scale, and the underlying factors contributing to this phenomenon, such as synergistic interactions, remain largely unexplored. The possibility remains, notwithstanding all the adaptive explanations, that defence islands evolve neutrally through a preferential attachment process whereby multiple defence systems are incorporated into genomic regions devoid of essential genes where insertions are tolerated.

Here we report a comprehensive analysis of the co-occurrence of defence systems in 26,362 *Escherichia coli* genomes, as well as in complete genomes from four bacterial orders, Enterobacterales, Bacillales, Burkholderiales, and Pseudomonadales, to investigate the role of interactions between different defence systems in anti-phage immunity. Our findings show that defence system co-occurrence varies substantially across *E. coli* phylogroups and taxa and is not directly related to their co-localisation in the genome. For several pairs of non-randomly co-occurring and mutually exclusive defence systems in *E. coli*, we experimentally demonstrated synergistic interactions that provided an evolutionary advantage to the bacterial population. Moreover, some of the defence systems that are mutually exclusive in *E. coli* were found to co-occur in other bacterial taxa, and can also protect synergistically against specific phages. These findings imply that selection for robust immunity, rather than mechanistic incompatibility, is the primary driving force that shapes the defence system repertoire in bacteria.

## RESULTS

### Distinct defence system repertoires across *E. coli* phylogroups

To explore the variation in the immune repertoires among closely related bacteria, we analysed the defence system content in a comprehensive dataset of 26,362 *E. coli* genomes from the NCBI Reference Sequence (RefSeq) database^35,36^. *E. coli* is an ideal model organism for this research due to its wide distribution in diverse environments, high genetic diversity, the availability of numerous, well-characterised, complete genomes as well as a large panel of well-studied phages^37,38^. We found that, in agreement with previous observations, on average, *E. coli* genomes carry 5-7 defence systems, but some clades, such as those in phylogroup B2.1, harbour a greater diversity of such systems (**Figure S1A**, **Table S1**). The majority of the defence systems are encoded in the chromosomes, but there are some clades where additional systems, especially Gabija, tmn, PifA, ppl, AbiQ, and viperins, are carried on plasmids (**Figure S1B**).

We sought to investigate in greater detail the clade-specific patterns and the differences among the six major *E. coli* phylogroups that vary in their ecology. Our dataset prominently represented phylogroups A (25%), B1 (29%), and B2 (19%), which are highly prevalent in the human (A, B2) or domestic/wild animal microbiomes (B1)^39^ (**Figure 1A**). Our analysis revealed substantial differences in the number (**Figure 1B**) and types (**Figure 1C**) of defence systems among the phylogroups. Phylogroup B2.1, which includes extra-intestinal pathogenic (ExPEC) strains^40^, was particularly noteworthy, with the highest average number of defence systems (8) among the examined phylogroups (**Figure 1B**). The genomes in phylogroup B2 accumulate virulence factors^41,42^ as well as antibiotic resistance genes^43^, suggesting that this phylogroup is specifically prone to HGT mediated by MGEs such as pathogenicity islands and plasmids. Of particular interest in phylogroup B2 is the already reported absence of CRISPR-Cas type I-E that is common in other *E. coli* phylogroups^44^ (**Figure 1C**). Conversely, significant enrichment (Chi Squared test for homogeneity, p < 0.001, **Table S2**) of Retron I-C (odds ratio (OR) = 9.04) and AbiE (OR = 4.77) was detected in the B2-1 subgroup, and high prevalence of CRISPR-Cas type I-F (OR = 5.12), Thoeris I (OR = 6.16), Septu I (OR = 3.17), and qatABCD (OR = 4.63) was observed in the B2-2 subgroup.

**Figure 1.**
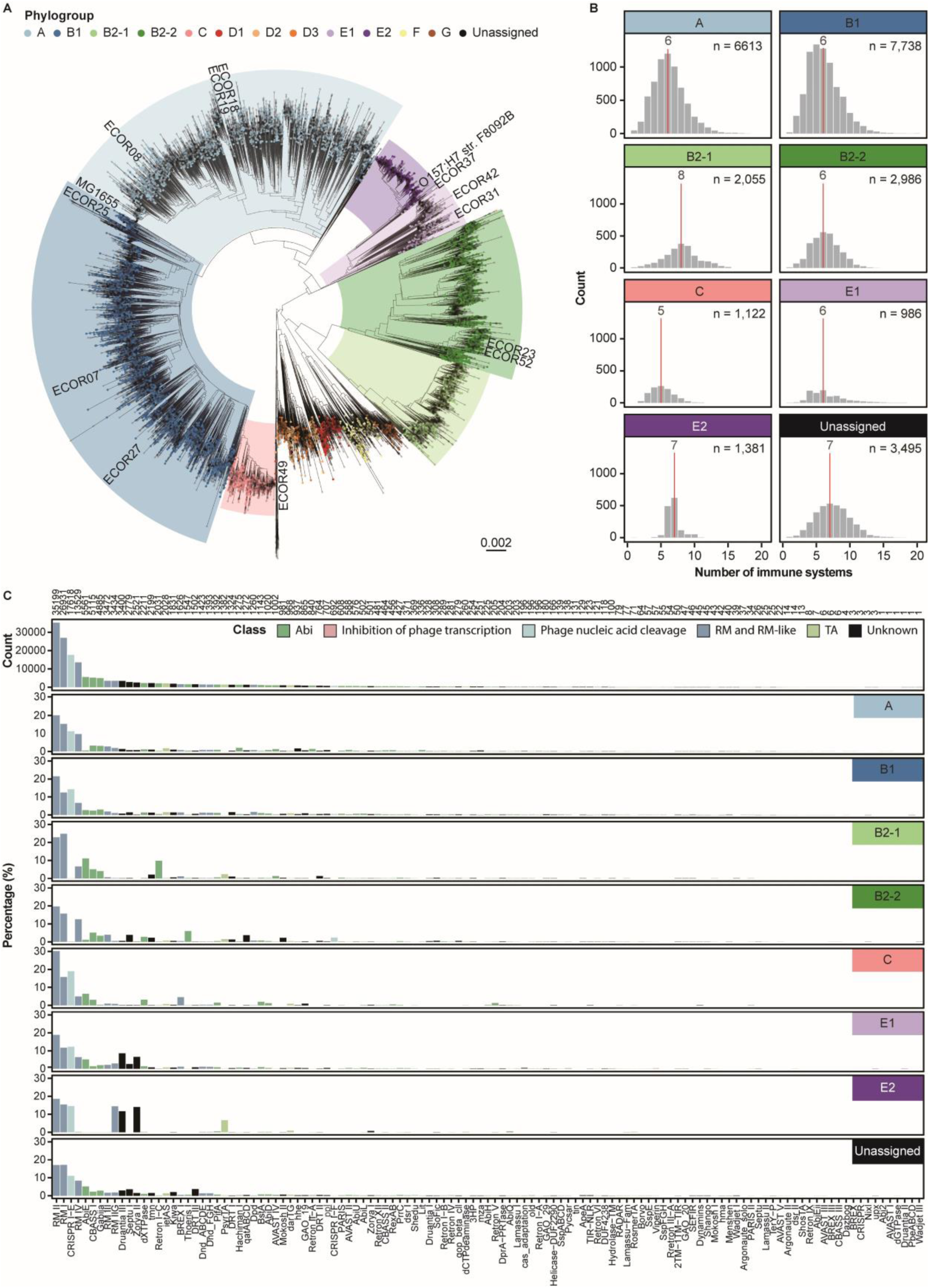
Distribution of defence systems across *E. coli* phylogroups. **(A)** A phylogenetic tree displaying 26,362 *E. coli* genomes obtained from the RefSeq database. Phylogroups are assigned as described previously^45^, and each phylogroup is colour-coded according to the key. **(B)** Number of defence systems found per *E. coli* strain in each phylogroup. The mean number of defence systems is indicated by a red line. **(C)** Prevalence of defence systems in the *E. coli* genomes. The defence systems are organised from most prevalent (left) to least prevalent (right) and their total count is shown in the top bar graph. The bars are colour-coded according to the mechanism of the defence system. The remaining bar graphs show the prevalence, in percentage, of the defence systems per phylogroup.

Phylogroup C showed enrichment of BREX I (OR = 3.54), and phylogroups E1 and E2 exhibited a much higher prevalence of Zorya II (OR = 4.62 and 10.25, respectively) and Druantia III (OR = 4.86 and 6.92, respectively) compared to the other phylogroups. Phylogroup E2 additionally showed a higher prevalence of PsyrTA (OR = 7.81), and a reduced prevalence of RM IV (OR = 0.02) that seems to be compensated by an increase in RM IIG (OR = 7.86).

For most phylogroups, we observed a relatively strong positive correlation (r = 0.34-0.66, p < 2.2x10^-16^) between the types of defence systems found in *E. coli* genomes within phylogroups and the genetic relatedness of these genomes, with the exception of phylogroups A and B1 (r = 0.02-0.11, p < 2.2x10^-16^), as indicated by the mash distance analysis (**Figure S1C**). These observations indicate that, although HGT is an important route of defence system acquisition, vertical inheritance plays a major role in the evolution of the immune repertoires, at least at short phylogenetic distances. The low correlation in phylogroups A and B1 likely reflects ecological differences between subclades in which case HGT apparently becomes a defining factor.

In summary, our analysis of the distribution of defence systems across *E. coli* genomes demonstrates associations between specific defence systems and individual phylogroups, likely driven by selection for sets of defence mechanisms capable of efficiently protecting the bacteria against the specific repertoires of phages and other MGEs that they encounter in their respective environments.

### 118 pairs of defence systems co-occur in *E. coli*

In previous studies, some defence systems have been shown to interact, resulting in enhanced or expanded protection against phages^29,31–34^. Here, we sought to determine if specific defence systems co-occurred more frequently than expected in bacterial genomes, potentially indicating interactions between different defence mechanisms enhancing protection against phages. To this end, we explored correlations between the occurrences of pairs of defence systems in *E. coli* genomes corrected for phylogenetic bias; we deemed such a correction to be essential because, as shown above, vertical inheritance of defence systems is common (see Methods). This analysis allowed us to identify pairs of defence systems that appear together in the same genome significantly more (co-occurring) or less (mutually exclusive) frequently than expected based on their individual prevalence (**Table S3**, **Figure 2A**).

**Figure 2.**
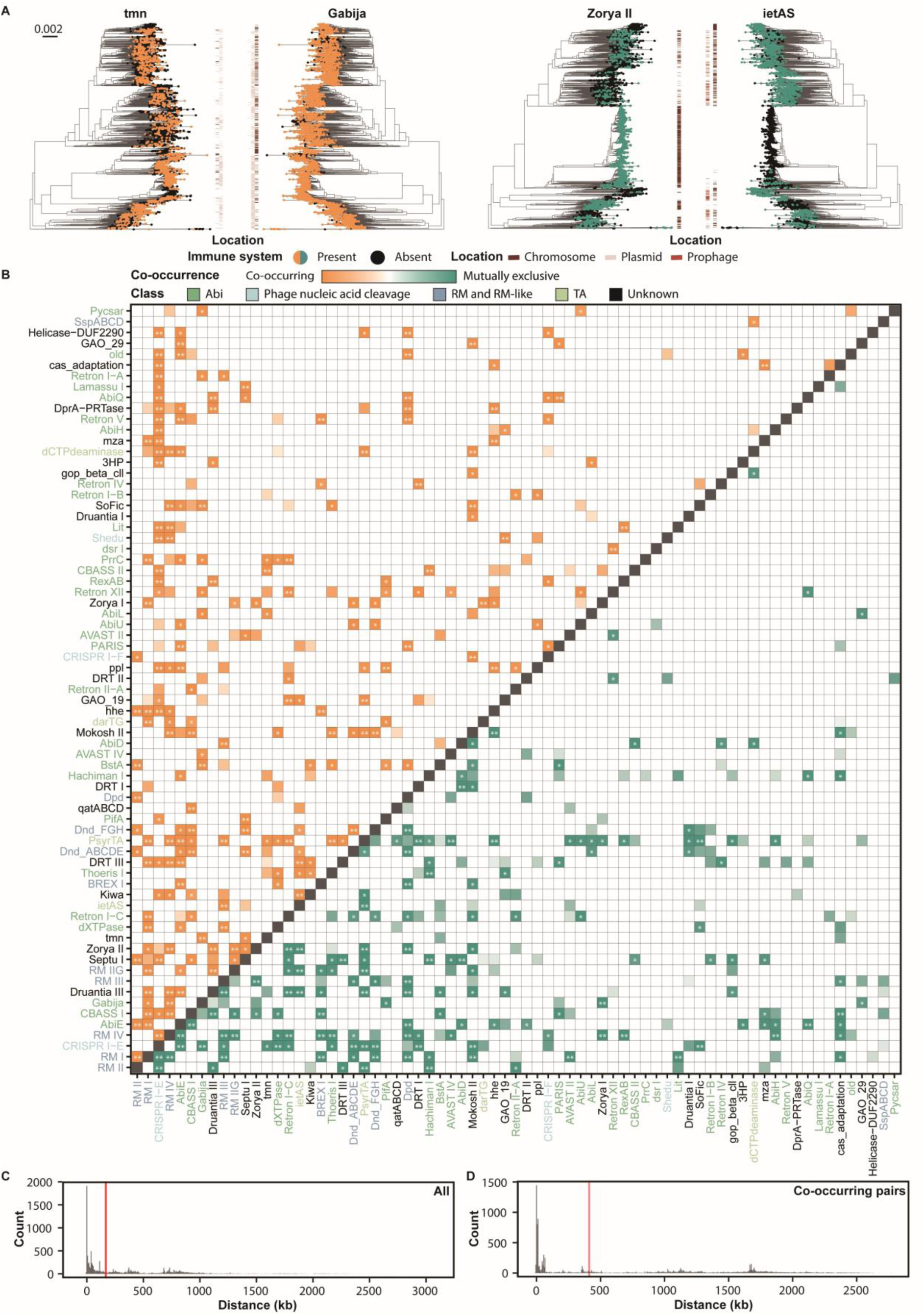
Co-occurrence and mutual exclusion among defence systems in *E. coli*. **(A)** Graphical representation of the co-occurrence analysis, depicting one pair of defence systems that co-occur (Gabija and tmn), and one pair that is mutually exclusive (Zorya II and ietAS). The nodes of the *E. coli* phylogenetic tree are coloured according to the presence or absence of the defence system in each strain. Their location in chromosome, plasmid, or prophage regions is indicated in the middle. For visualization purposes, only leaves that carry at least one system from the pair are shown. **(B)** Co-occurrence of defence system pairs in *E. coli*. Co-occurring systems are shown in orange, and mutually exclusive systems are shown in green. The correlation between pairs was calculated with Pagel test for binary traits. Asterisks show correlations that were significant after the less stringent Benjamini-Hochberg correction (*) or the most stringent Bonferroni correction (**) for multiple testing. In the main text, the results after Bonferroni correction are considered. The defence systems are colour-coded according to their broadly defined mechanism of defence. **(C,D)** Distance histograms of (B) all and (C) co-occurring defence system pairs in 2,164 complete *E. coli* genomes. The median distance between genes encoding defence systems is shown by a red line. The analysis considered only those pairs that significantly co-occurred after Bonferroni correction.

Our analysis revealed that 183 interacting pairs of defence systems (7.4% of all the analysed pairs) were significantly correlated, positively or negatively (Pagel test for binary traits, with Bonferroni correction for multiple tests). Of these, 118 pairs were co-occurrences (64.5% of the correlated pairs), and the rest were cases of mutual exclusion. With the more permissive Benjamini-Hochberg correction, 379 pairs of defence systems (15.3% of the analysed pairs) were significantly correlated, of which 219 (57.8%) were co-occurrences (**Figure 2B**). Notably, although the network of co-occurrences and especially of exclusions between the *E. coli* defence systems was sparse (**Figure 2**, **Figure S2A**), each system significantly co-occurred with at least one other system, and typically, with two or more under the permissive correction, and for most, at least one significant co-occurrence was detected under the strict correction, too (**Table S3**). Thus, co-occurrence between defence systems is a widespread phenomenon that involves (nearly) all such systems identified in *E. coli*. The greatest number of significant co-occurrences was observed for the CRISPR I-E system that is found in the majority of the *E. coli* genomes, but several less common systems, such as AbiE, PsyrTA and Mokosh II, also appeared to be particularly prone to co-occurrence with other systems (**Table S3**). Some of the co-occurring pairs appeared striking in that the great majority of the instantiations of the rarer system in the pair were found in genomes that also carried the more common system, suggestive of a functional dependence and indeed their interaction was better described by the dependent model (**Table S3**). For example, 882 of the 975 instances of Mokosh II co-occurred with RM IV, and 2151 of the 2518 instances of Zorya II co-occurred with Druantia III (**Figure 2A** and **Table S3**).

Notably, different subtypes of the same defence system displayed distinct co-occurrence patterns with other systems (**Figure 2B**, **Figure S2B**). For example, while Druantia III co-occurred with 8 other defence systems, no co-occurrences were found for Druantia I (Bonferroni correction). CBASS I, composed of cyclase and effector proteins, and CBASS II, characterised by the presence of cGasylation proteins cap2 (E1-E2 fusion) and cap3 (JAB)^19,46^, co-occurred with distinct defence systems. CBASS I co-occurred with DndABCD and DndFGH (which also co-occurred with each other), Mokosh II, qatABCD, and RM I and IV, whereas CBASS II co-occurred with CRISPR-Cas type I-E, Hachiman I, and tmn. These specific co-occurrences might reflect distinct cooperative interactions between the respective defence system subtypes.

Conversely, we observed 65 (2.6%) pairs of defence systems that were mutually exclusive, such as CRISPR-Cas types I-E and I-F1, consistent with previous observations in Proteobacteria^31^ (**Figure 2B**). Like the number of co-occurrences, the number of significant exclusions notably varied across defence systems, with some, for example, PsyrTA and RM IV appearing particularly prone to avoiding other systems (**Figure S2B**). Interestingly, RM IV and BREX I were mutually exclusive in *E. coli*, even though previously shown to co-occur on a plasmid-encoded defence island and provide complementary protections against modified (RM IV) and non-modified (BREX I) invading DNA in *Escherichia fergusonii*^29^. However, here we consolidated all subtypes of RM IV together, and the majority of occurrences in our dataset were in the chromosome, showing that the observed pattern in *E. fergusonii* was specific and differed from the typical behaviour of RM IV. Additionally, location of one of the systems within an integrated element, such as prophage, can lead to mutual exclusion with systems active against that MGE, as it potentially happens in the case of ietAS and Zorya II (**Figure 2A**). The RM-like Dpd system, which acts by inserting 7-deazaguanine derivatives into the host DNA to distinguish it from the non-modified invading DNA^47^, was found to be mutually exclusive with several RM and RM-like systems, such as RM I, RM III, RM IV, BREX I^48–50^, Druantia III^2^, and DndABCD and DndFGH^9^.

Like the co-occurrence patterns, the patterns of mutual exclusion showed substantial differences among subtypes of the same defence system (**Figure 2B**, **Figure S2B**). In most cases, however, the exclusivity between defence systems was not strict, that is, the respective pairs were observed together in some genomes (**Table S3**). This observation implies that the exclusivity is not caused by incompatibility between the respective systems resulting in negative epistasis^31^, but rather by genetic drift due to functional redundancy or by selection against such redundancy.

Overall, our results indicate that both non-random co-occurrence and (partial) mutual exclusion among defence systems are common in *E. coli*.

### Co-occurrence of defence systems is not tightly linked to physical proximity

Defence systems often cluster together in defence islands^2,4,7,24,26,51^, which has been hypothetically attributed to fitness benefits conferred by such clustering on bacteria living in environments with high phage loads ^27^. In particular, clustering of defence systems in defence islands, especially within integrated MGEs, increases the likelihood of horizontal co-transfer^52^. Therefore, if the defence system repertoire is predominantly shaped by HGT, and not by any functional benefits, the co-localizing systems will also be the ones that we observed as co-occurring in *E. coli*. However, our analysis showed that co-occurring defence systems were generally not located significantly closer to each other in the genome compared to the average distance between defence systems (**Figure 2C,D**). Thus, in general, the physical proximity of defence systems within the genome, although likely facilitating their concomitant horizontal transfer, does not appear to play a defining role in their co-occurrence.

Nonetheless, there are some exceptions where co-occurring defence systems did indeed non-randomly co-localize. These include Mokosh II and RM IV, Druantia III and RM I, Druantia III and RM IV, RM I and RM IIG, RM IIG and Zorya II, Druantia III and Zorya II, Druantia III and RM IIG, Gabija and tmn, DndABCDE and DndFGH, CBASS I and qatABCD, RM III and ietAS, and RM IV and SoFic (**Figure S2C**). Of these pairs, only DndABCDE and DndFGH were previously reported to co-localize and functionally interact^53,54^.

### Co-occurring defence systems act synergistically to counter phage infection

Next, we sought to explore whether co-occurrence of defence systems is driven by their complementary activities^30^ or synergistic molecular cooperation^28,29^. To this end, we selected three pairs of significantly co-occurring defence systems found in *E. coli* strains from our collection: Gabija and tmn that co-localise frequently in plasmids, Druantia III and Zorya II that co-localise in the chromosome, and ietAS and Kiwa that do not co-localise (**Figure S2C**). Additionally, we tested the mutually exclusive combination of Zorya II and ietAS.

By cloning these systems into *E. coli* individually or jointly (**Figure 3A**), we assessed their effects on phage resistance using efficiency of plating (EOP) assays with a panel of 29 phages. This experiment demonstrated limited anti-phage activity for all single systems, with the exception of Gabija, which provided strong protection against multiple phages (**Figure S3A**). Combinations of the defence systems substantially increased the protection levels against specific phages. To further quantify these effects, we calculated the epistatic coefficients by comparing the combined effect of the defence system pair to the sum of the individual effects of the two partners. The results showed that all tested combinations of defence systems, with the exception of ietAS and Kiwa, displayed significant synergistic effects against at least some phages in our panel, with even the mutually exclusive pair Zorya II and ietAS showing unexpected synergy (**Figure 3B**, **Figure S3A**).

**Figure 3.**
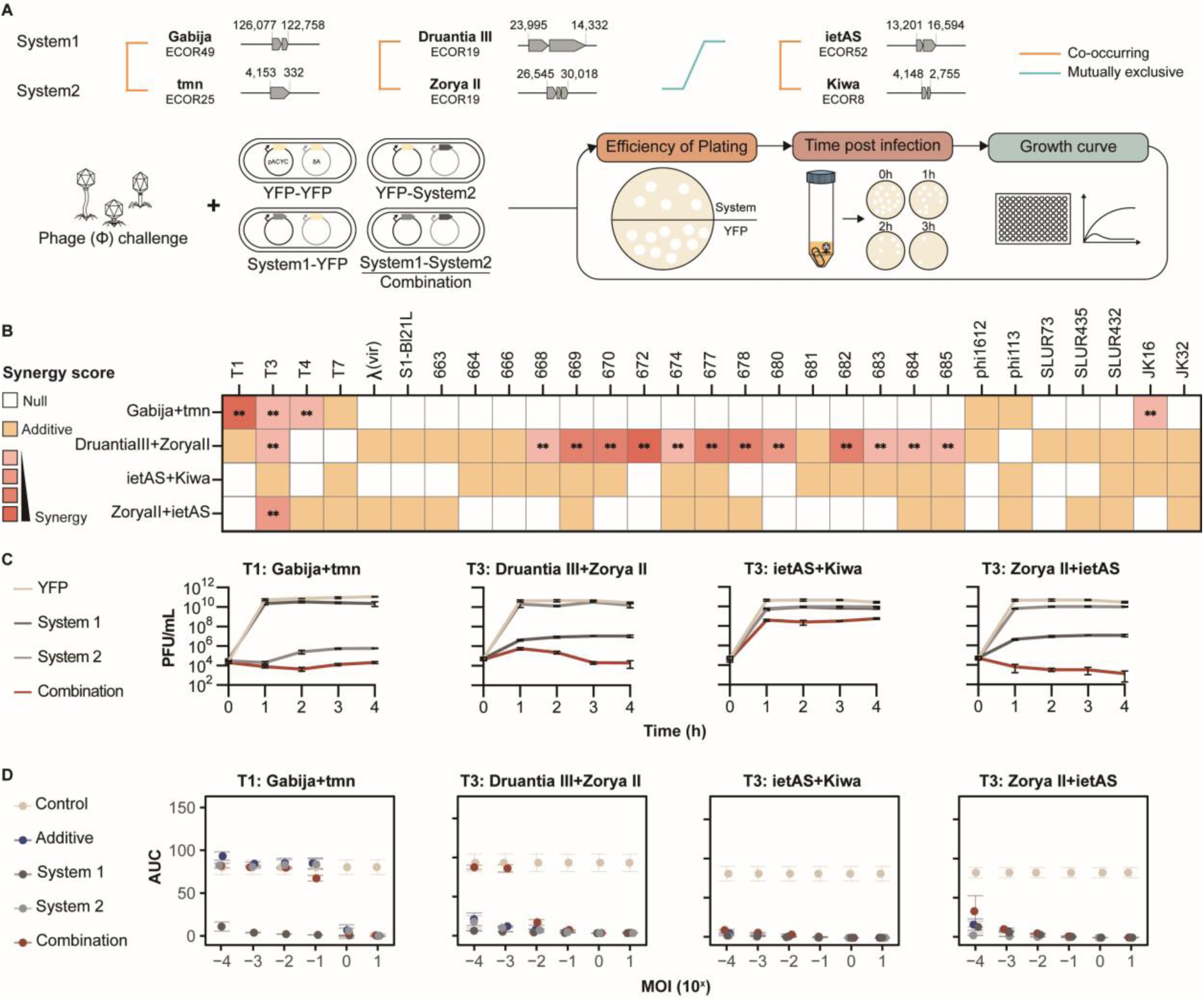
Defence system pairs provide synergistic anti-phage activity. (A) Experimental set-up for the assessment of the anti-phage activity of individual defence systems and their combinations. YFP, yellow fluorescent protein. (B) Heatmap of synergy score of protection provided by selected defence system pairs against a panel of 29 phages. The synergy score is the epistatic coefficient for pairs of defence systems (see Methods). Null, EOP equivalent to the defence provide by one system; Additive, EOP corresponds to the combined defence of the two individual systems; Synergy, EOP exceeds the collective defence of the two systems. Data is shown as the average of three biological replicates. ** Statistically significant (p < 0.01). (C) Time post infection assays, measuring T1 or T3 titres over the course of four hours in liquid cultures of *E. coli* containing individual or combined defence systems. Data is shown as the average and standard deviation of three biological replicates. (D) Bacterial growth under phage predation at different multiplicities of infection (MOIs), represented as area under the curve (AUC) in OD⋅h. An defence system pair acts synergetic when its dot (red) is above the expected additive effect (blue). Data is shown as the confidence interval of three biological replicates. The raw data and growth curves used to calculate the AUCs are available on the associated Github and Zenodo databases.

To further validate the findings from the EOP assays, we performed time post-infection assays using phages T1 and T3 to assess the impact of the defence system combinations on phage propagation in liquid cultures. The results from these assays consistently confirmed the synergy as the combinations of defence systems led to a reduction in phage propagation that significantly exceeded the sum of the individual effects (**Figure 3C**, **Figure S3B**). This synergistic trend was observed for phage T1 with Gabija and tmn, for T1 and T3 with Druantia III and Zorya II, and for T3 with Zorya II and ietAS. Even the combination of ietAS and Kiwa, which showed no significant synergy in the EOP assays (**Figure 3B**), displayed a minimal but detectable synergy against T3 propagation (**Figure 3C**). Moreover, the synergy was not restricted to phages targeted by both individual systems. For example, T1 was not affected by Gabija alone, but the combination of Gabija and tmn resulted in an increased protection compared to tmn alone (**Figure 3C**). For the mutually exclusive pair Zorya II and ietAS, only Zorya showed substantial activity against phage T3, but the combination with ietAS resulted in an improved reduction in phage propagation.

Additionally, we assessed synergy between defence systems in terms of bacterial survival by measuring the absorbance of bacterial cultures over time when infected with phages at different multiplicities of infection (MOIs) (**Figure 3D**). To quantify the synergy in these assays, we compared the areas under the curve (AUC) above the OD at the experiment start for individual systems and their combinations. These additional results from liquid cultures were consistent with the findings from the EOP and phage propagation assays, providing further evidence of synergistic interactions between the defence systems. We observed synergy occurring at low MOIs, resulting in increased bacterial survival. However, at higher MOIs, all bacterial cultures collapsed, likely due to overwhelming of the defence systems or abortive infection. In the case of Gabija and tmn, increased bacterial survival was not observed at low MOIs for the combination of systems because tmn alone was sufficient to restore normal growth of T1-infected bacteria (**Figure 3D**, **Figure S3C**).

Taken together, our assays provided robust evidence that some of the significantly co-occurring defence systems display synergistic activity against specific phages. This bolsters the notion that co-occurrence is maintained by environmental selection favouring combinations that are beneficial for the bacteria given the particular virome composition in their niche. Additionally, the observation that mutually exclusive defence systems in some cases also display synergistic activity suggests that inherent mechanistic incompatibilities between defence systems are unlikely to be a major driver of their mutual avoidance.

### Defence system co-occurrence is not conserved across bacterial taxa

Genomic analyses show that increasing phylogenetic distance between bacterial species, decreases the rate of HGT^55–58^. In particular, the host ranges of most phages are narrow so that transduction occurs mostly within the host species boundaries^59^. Similarly, only a small fraction of plasmids has been shown to cross the interspecies barrier^60–62^. Considering these limitations of HGT along with (largely) non-overlapping viromes^22^, it could be expected that the co-occurrence of defence systems is poorly conserved among bacteria. To test this prediction, we extended our analysis to four bacterial orders, Bacillales, Burkholderiales, Enterobacterales, and Pseudomonadales. We first examined the sets of defence systems present in these bacteria. The different bacterial orders displayed variations in the type and abundance of defence systems (**Figure S4A**). For example, RM systems were the most prevalent in Bacillales, Enterobacterales (including *E. coli*), and Pseudomonadales, whereas Burkholderiales were characterised by a higher abundance of dXTPase, Zorya III, and Mokosh I. Additionally, while CRISPR I-E was abundant (50.8%) in Enterobacterales (including *E. coli*), its prevalence was markedly lower (<6%) in the other orders. Nonetheless, when considering all four orders and one extensively characterized species (*E. coli*), they collectively shared 87 defence systems in common, with only a few systems unique to each of them (3 in *E. coli*, 18 in Bacillales, 2 in Enterobacterales, 2 in Pseudomonadales, and 1 in Burkholderiales) (**Figure S4B**).

As anticipated, the specific pairs of co-occurring and mutually exclusive defence systems differed across the taxa. Moreover, multiple pairs of defence systems that co-occurred in one order were found to be mutually exclusive in another (**Figure 4**). For instance, AbiE and RM I co-occur in Bacillales and Enterobacterales but are mutually exclusive in Burkholderiales. Similarly, CBASS type II and CRISPR-Cas type I-E co-occur in Enterobacterales but are mutually exclusive in Pseudomonadales. These findings indicate that mutually exclusive systems are not inherently functionally incompatible or redundant. Rather, the interactions between these systems likely depend on environmental and genetic factors that select for a particular anti-phage immunity strategy. Overall, our results indicate that defence systems are generally mechanistically compatible, allowing bacteria to adopt diverse, flexible strategies for anti-phage defence based on their unique environmental and genetic contexts.

**Figure 4.**
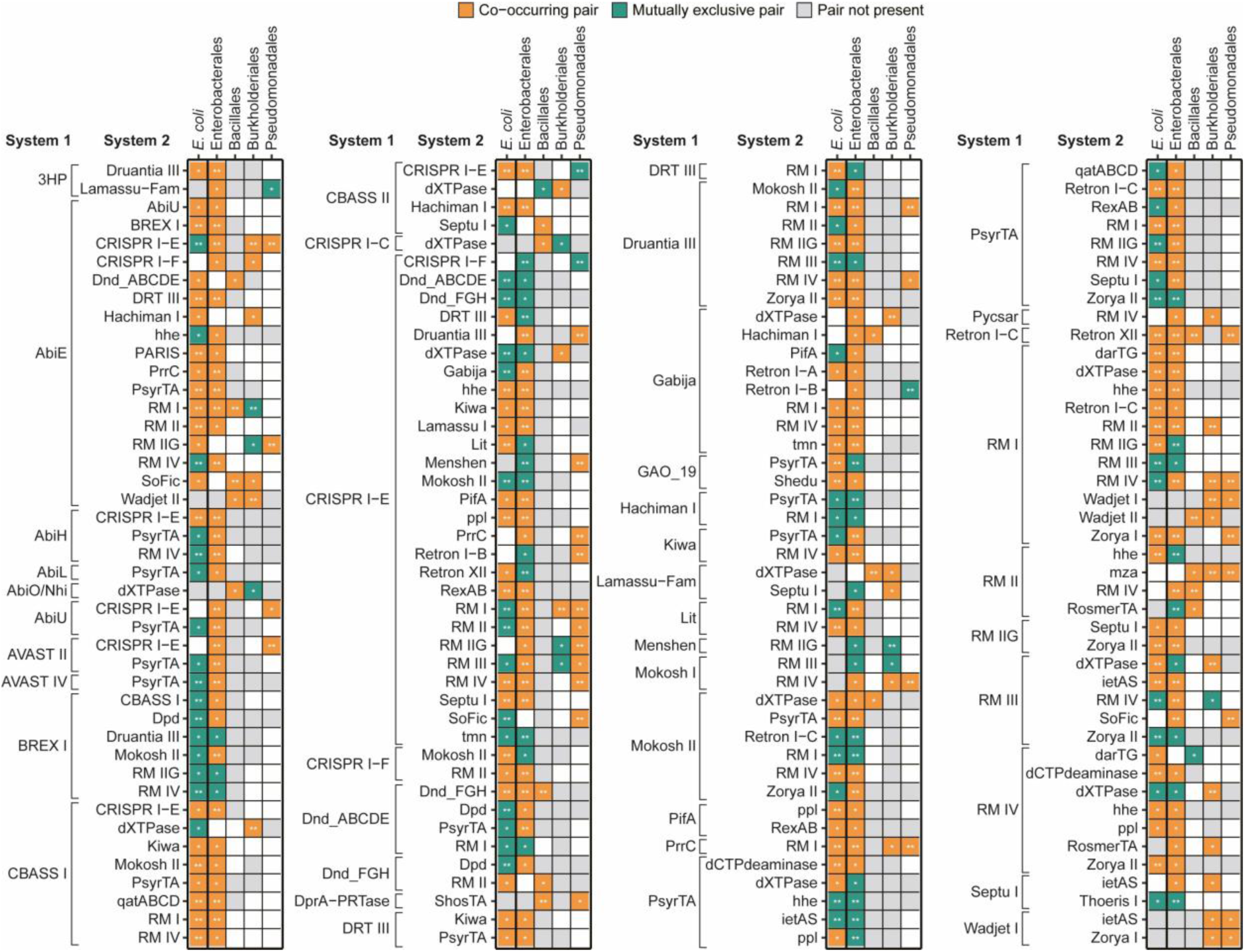
Patterns of defence system co-occurrences across bacterial taxa. Heatmap of defence system co-occurrence patterns in *E. coli* (n = 26,362) and in four bacterial orders: Enterobacterales including *E. coli* (n = 9,124), Bacillales (n = 3,952), Burkholderiales (n = 2,199), and Pseudomonales (n = 1,288). Grey squares indicate that at least one system in the pair is not present in the taxonomic group. * Co-occurrence significant after Benjamini-Hochberg correction; ** Co-occurrence significant after Bonferroni correction.

### Synergistic immunity provides an evolutionary advantage to bacterial populations

Due to our observations, we reasoned that co-occurring and synergistic defence systems could provide advantages at the population level. To assess the evolutionary and ecological impact of synergistic interactions between defence systems, we performed a short-term evolution experiment using chromogenic reporter plasmids expressing engineered coral chromoproteins^63,64^. We mixed populations of strains containing either an individual defence system and a second, “empty” chromogenic plasmid, or carrying two defence systems on separate plasmids. These populations were then infected with either a phage shown to trigger a synergistic defence, or a phage eliciting no obvious synergy between the respective defence systems. Over a period of three days, we monitored the proportion of the population carrying one or both defence systems by counting colony forming units of different colours (**Figure 5A**). We confirmed that all plasmids were stably maintained in the populations throughout the experiment using plasmid loss assays, ruling out any influence of plasmid loss due to toxicity on the outcomes (**Figure S5A,B**). Further, given that resistant receptor mutants tend to spread in bacterial populations shortly after phage infection^65^, we investigated whether this factor influenced the outcome of our short-term evolution experiments by evaluating the capacity of the phage to infect bacterial colonies retrieved from the experiment. Since the defence systems under examination in this study do not offer full protection against phage infection, the absence of infection indicates the emergence of receptor mutants leading to complete phage resistance. We observed a limited effect of receptor mutants on phage resistance, with greater relevance at high phage MOI (**Figure 5B**).

**Figure 5.**
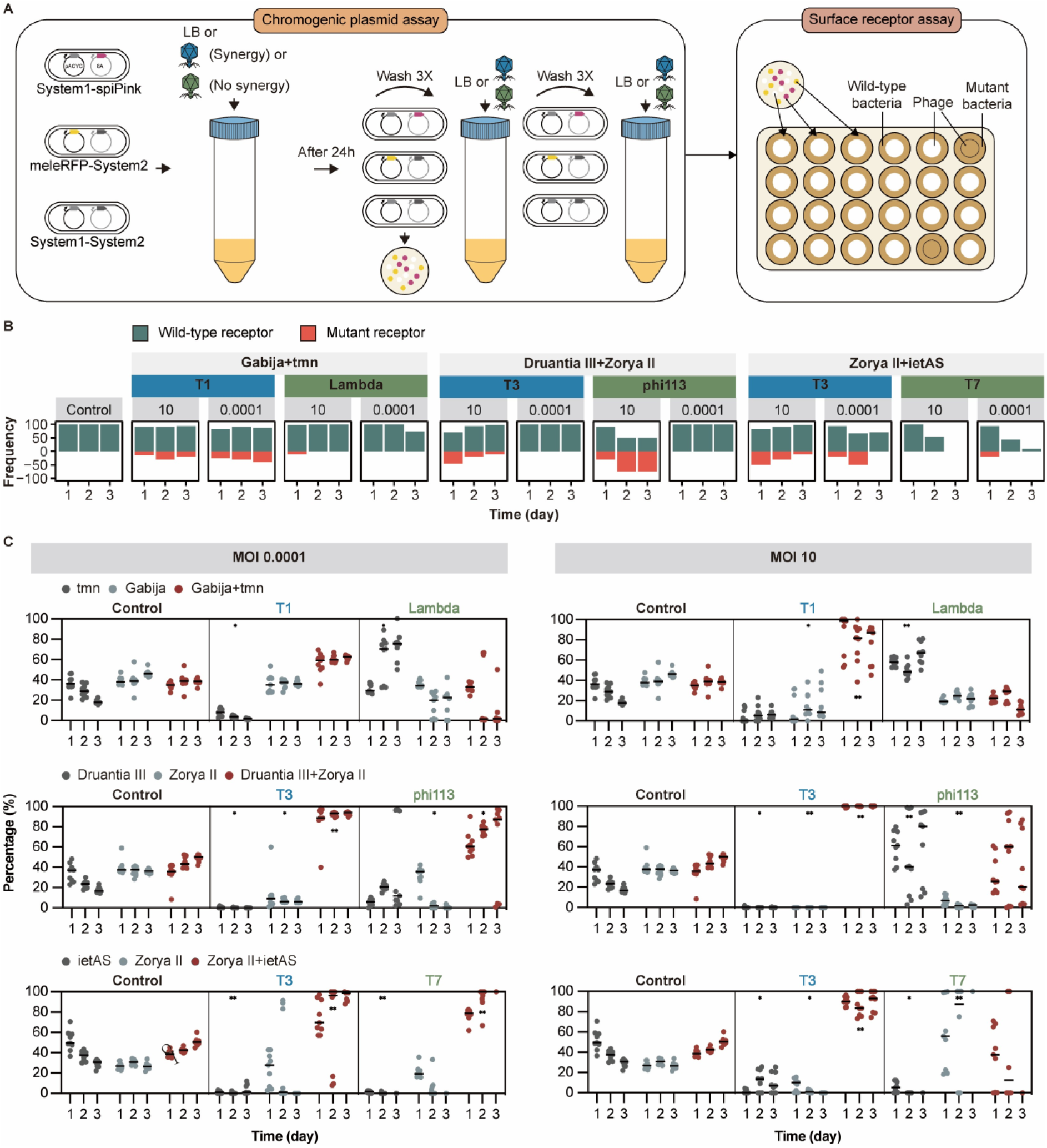
Synergistic defence system pairs provide an evolutionary advantage to bacteria. **(A)** Set-up of experimental evolution assay of defence systems using chromogenic plasmids (left). Cells containing a single defence system carry a second plasmid that expresses a chromogenic reporter. Cells with defence system combinations were mixed in equal proportions with cells with single systems, and infected with phage. After 1, 2, and 3 days, cells were platted and the colonies of different colours were enumerated. A surface receptor assay (right) assessed the influence of receptor mutants on the outcome of the evolution assays, by subjecting colonies of different colours to phage infection in plaque assays. **(B)** Prevalence of receptor mutants in the bacterial population during the evolution assay. The proportion of colonies with the wild-type receptor is represented as positive in the vertical axis, while the proportion of colonies with mutated receptor is shown as negative. Colonies with mutated receptor were identified by their complete resistance to phage infection in spot assays. **(C)** Percentage of colonies from the evolution assay that carry each individual defence system or their combinations at 1, 2 and 3 days post infection with phage at low or high multiplicity of infection (MOI), compared to a non-infected control. Data is shown as the average of three biological and three technical replicates with individual counts shown. * *p* value < 0.01, ** *p* value < 0.001 determined by multiple comparison for nested one-way ANOVA, comparing to the corresponding uninfected control.

The results showed that in the absence of phage, populations containing either individual or combined defence systems remained relatively stable, and thus these systems were minimally toxic to the cells, if at all (**Figure 5C**). However, after exposure to phage infection at high or low MOI, a clear shift in the population composition towards cells harbouring both defence systems was observed starting from day 1. This shift was pronounced in cultures infected with the phage that elicited synergistic defence (T1 for Gabija and tmn, T3 for Druantia III and Zorya II, and T3 for ietAS and Zorya II). In cases the defence was not synergistic (Lambda for Gabija and tmn, phi113 for Druantia III and Zorya II, and T7 for ietAS and Zorya II), the outcomes varied. For Druantia III and Zorya II, as well as Zorya II and ietAS combinations, cells carrying both the combination of systems and the individual system active against the phage (Druantia III and Zorya II, respectively **Figure S3A**) became predominant in the population. However, in the case of the Gabija and tmn combination, the system that was not active against the phage (tmn) dominated (**Figure S3A**). This discrepancy can be attributed to the considerably higher efficiency of Gabija against phage Lambda (10^5^-fold) compared to that of Druantia III against phi113 (10^1^-fold) and Zorya II against T7 (10^1^-fold) (**Figure S3A**). This result suggests that the phage population was more effectively reduced when Gabija was active, allowing other cells in the population, even those lacking active defences against the phage, to survive. In the case of the Gabija-tmn combination, the reduction in the population of cells containing the active system, Gabija, was likely due to the abortive infection phenotype of this defence system^66^.

These results underscore the pivotal role played by the degree of protection provided by specific, active sets of defence systems in the survival of phage-sensitive cells within a heterogeneous population, thereby shaping the dynamics of coexisting defence systems. Overall, the competition experiments validate the evolutionary advantage of synergistic defence system combinations against specific phages. Moreover, these findings also emphasise the substantial impact of factors such as the type and abundance of the encountered phages, as well as the effectiveness of the defence systems, on the resulting dynamics of the bacterial population.

### Tmn co-opts the ATPase domain of Gabija for synergy

To better understand the molecular mechanisms underlying the observed synergistic effect between defence systems, we focused on the combination of Gabija and tmn. This choice was motivated by several factors. Gabija has been well characterised previously, in contrast to all other tested defence systems, providing molecular detail^2,66–68^ that helps understanding the synergy with the less thoroughly characterized tmn^3^. Furthermore, Gabija and tmn tend to physically co-localise on plasmids (**Figure S2D**). Comparison of plasmids carrying both tmn and Gabija showed pronounced genetic variability, except for regions that encompass conjugation-related genes and these particular defence systems (**Figure 6A**). Gabija and tmn, along with type II TA systems such as VapBC, are specifically located in the leading region of the plasmid. This region is crucial for maintaining plasmid stability during conjugation^69,70^, and is enriched in anti-defence genes^71^, which protect conjugative plasmids from host defence systems during the initial stages of plasmid invasion. The conserved location of Gabija and tmn within the leading region of plasmids suggests that they play a critical role in plasmid maintenance. By allowing the plasmid to fend off other, competing MGEs, these systems likely ensure the plasmid maintenance in the cell, and with it, the evolutionary success of this defence system combination in the face of ongoing inter-MGE conflicts^28^.

**Figure 6.**
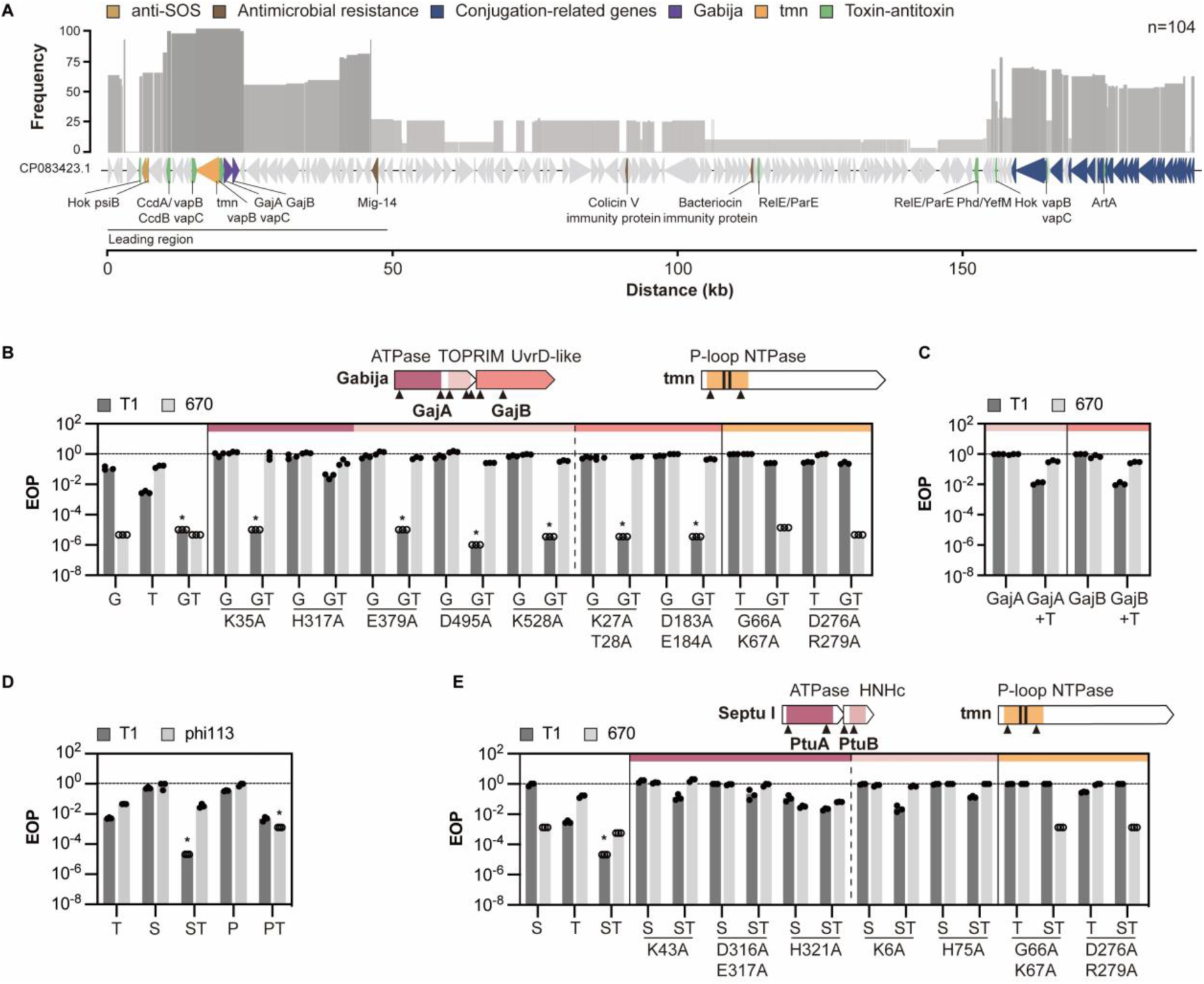
Mechanistic insight into the synergistic interaction between tmn and Gabija or Septu I. **(A)** Whole-plasmid alignment of 104 plasmids containing tmn and Gabija from complete *E. coli* genomes with plasmid CP083423.1 as a reference (see Methods). The histogram shows the percentage of plasmids where the corresponding block from CP083423.1 was found. Annotated genes are coloured by function. **(B)** Efficiency of plating (EOP) of phages T1 and 670 on cells expressing Gabija (G), tmn (T), Gabija and tmn (GT), and alanine mutants of specific functional domains. The mutations are organised by functional domains of Gabija (ATPase, TOPRIM, and UvrD-like) and tmn (P-loop NTPase). Unfilled circles indicate instances where it was not possible to determine the number of phage plaques, hence a value of 1 was assumed at the corresponding dilution. Asterisk (*) indicates cases of synergy. **(C)** EOP of phages T1 and 670 on cells expressing tmn (T), Gabija (G), and tmn with either GajA or GajB. Unfilled circles indicate instances where it was not possible to determine the number of phage plaques, hence a value of 1 was assumed at the corresponding dilution. **(D)** EOP of phages T1 and phi113 on cells expressing tmn (T), Septu I (S), PrrC (P), Septu I and tmn (ST), or PrrC and tmn (PT). Unfilled circles indicate instances where it was not possible to determine the number of phage plaques, hence a value of 1 was assumed at the corresponding dilution. Asterisk (*) indicates cases of synergy. **(E)** EOP of phages T1 and 670 on cells expressing tmn (T), Septu (S), and Septu and tmn (ST), and variants with point mutations in specific functional domains. The mutations are organised by functional domains of Septu I (ATPase and HNHc) and tmn (P-loop NTPase). Unfilled circles indicate instances where it was not possible to determine the number of phage plaques, hence a value of 1 was assumed at the corresponding dilution. Asterisk (*) indicates cases of synergy.

To explore the mechanism of synergy between tmn and Gabija, we focused on determining the specific contributions of the functional domains of the individual defence systems to the synergistic anti-phage activity. We introduced point mutations into these functional domains and assessed their effects on the individual defence systems and their combination using EOP assays.

In tmn, mutations of conserved residues of the P-loop NTPase domain (G66A/K67A, and R276A/R279A) abolished protection by tmn and the synergy with Gabija against phage T1 (**Figure 6B**). In Gabija, the ABC ATPase domain of GajA senses the depletion of cellular nucleotides during phage infection, activating the DNA binding and nicking activity of its TOPRIM nuclease domain^68^. Introduction of nicks into the DNA activates nucleotide hydrolysis by the UvrD-like helicase domain of GajB, depleting essential nucleotides and leading to abortive infection^66^. Surprisingly, we observed that only the nucleotide-sensing ATPase domain of GajA appeared to be critical for the synergy with tmn (**Figure 6B**) because mutations of conserved residues in the respective catalytic sites^66^ of the TOPRIM and UvrD-like helicase domains abolished protection by Gabija on its own but had no effect on the synergy. The individual K35A and H317A mutations introduced into the ATPase domain of GajA abolished Gabija activity but only H317A abolished the synergy with tmn. The histidine residue H317 has been shown previously to play a role in inhibiting the nicking activity of GajA in the presence of ATP, suggesting that this histidine is critical for sensing the nucleotide levels in the cell^66^ and for the synergy with tmn.

Gabija forms a supramolecular complex composed of a GajA tetramer with two sets of GajB dimers docked on opposite sides^67^. While the helicase function of GajB and the TOPRIM activity of GajA do not appear to be required for the synergy with tmn, we sought to determine whether the intact complex was a critical factor for the molecular interactions leading to synergy. We introduced stop codons into each Gabija protein-coding sequence and assessed the impact of halted protein translation on the synergy with tmn. Our findings indicate that GajA alone could not synergise with tmn (**Figure 6C**), highlighting the necessity of forming an intact GajAB supramolecular complex for the synergy.

These findings demonstrate the critical role of the ATPase domains of both GajA and tmn in driving the synergy between these defence systems. More specifically, the ABC ATPase domain of GajA enhances the activity of tmn, emphasizing the central role of tmn in the synergistic combination. The molecular mechanism of tmn enhancement remains to be elucidated. Tmn is a KAP NTPase^3^, characterised by the presence of two transmembrane helices inserted into the P-loop NTPase domain, which anchor tmn in the membrane such that the P-loop domain is located on the intracellular side^72^. Because most P-loop ATPases are multimers (most commonly, hexamers^73^), we modelled the tmn dimer using Alphafold2 (pLDDT score 89.1) (**Figure S6A**), and observed that the residues situated between the two transmembrane helices (P177-P198) are likely located in the periplasmic region. This periplasmic loop might be involved in the recognition of phage components resulting in the activation of the NTPase. It has been proposed that the function of the KAP NTPase domain is the regulation of assembly/disassembly of other protein complexes that interact with the extended surfaces provided by the α-superhelical structural domains of tmn, in an NTP-dependent manner^72^. Hence, the ABC ATPase of Gabija might assist the NTPase domain of tmn in this regulation, increasing the downstream response, by a mechanism that remains to be elucidated.

Overall, our exploration of the synergistic interaction between tmn and Gabija revealed a specific molecular interplay between these systems, highlighting the pivotal role of the distinct ATPase domains of each of these defence systems.

### Synergy between tmn and defence systems containing sensory switch ATPases

Because tmn appears to be the main driver of the synergy with Gabija, we examined the domain architectures of the other defence systems that significantly co-occurred with tmn, namely, AbiL, PrrC, PsyrTA, Septu I, and CBASS II (**Figure 2A**). Except for CBASS II, all these systems, including Gabija, contain ATPase domains associated with effectors that likely cause DNA or RNA damage^2,74–76^, suggesting a potential shared mechanism underlying synergistic interactions with tmn. To test this hypothesis, we analysed the anti-phage defence provided by tmn when combined with the co-occurring systems Septu I and PrrC. As observed for tmn and Gabija, the combination of tmn with Septu I or PrrC demonstrated synergistic effects against specific phages (**Figure 6D**). The synergy between tmn and Septu I was consistently observed when using Septu I gene clusters from different *E. coli* strains (**Figure S6B**).

We further characterised the synergy between tmn and Septu I by introducing mutations into critical residues of Septu I. Mutations in the Walker A (K43A), Walker B (D316A/E317A), and D-loop (H321) regions of the ATPase domain of PtuA, as well as in the predicted Mg^2+^ binding site (K6A) and active site (H75A) of the HNHc nuclease domain of PtuB, resulted in the loss of synergy against phage T1, indicating the essential role of both proteins in the synergy (**Figure 6E**). Both PtuA and PtuB proteins act as toxins of the retron Ec78 defence system, and it was proposed that PtuA, similarly to GajA in Gabija, is activated by NTP depletion during phage infection^77^. This feature is also observed in the PrrC defence system, where the ATPase domain of PrrC is inhibited by ATP and GTP, alongside negative regulation by the RM system PrrI. The release of this inhibition, triggered by phages deploying anti-restriction peptides that inhibit the PrrI restriction enzyme, leads to GTP hydrolysis and activates the C-terminal anti-codon nuclease (ACNase) HEPN domain^75,78,79^. The PsyrTA^76^, also known as RqlHI^80^, and AbiL^74^ defence systems, which co-occur with tmn, likely function in a similar manner, given that the ATPase containing proteins of these systems have been shown to inhibit the toxic activity of the other protein.

In summary, our results show that tmn synergises with various defence systems containing ATPase domains that function as sensory switches unmasking the associated effector domains, as observed here with Gabija, PrrC, and Septu I. These domains likely aid the NTPase domain of tmn in controlling its downstream response. Further exploration of the mechanisms of these defence systems is expected to provide deeper insights into the synergistic interactions.

## DISCUSSION

In this work, we aimed to gain insights into the interactions between defence systems and the impacts of such interactions on bacterial immunity against phage predation both at cellular and population level. By comprehensive analysis of thousands *E. coli* genomes, we identified patterns of co-occurrence as well as mutual exclusion among defence systems. We showed that the co-occurrences are not conserved in more distant bacteria, suggesting the existence of many distinct defence strategies. Perhaps unexpectedly, the co-occurrence among defence systems was found not be strongly linked to their co-localisation in bacterial genomes, although we did identify several instances in which co-occurring systems co-localised. In several specific cases we explored, co-occurrence of both co-localising and non-co-localising pairs of systems was associated with synergistic interactions which led to a significantly greater protective effect against specific phages than expected from the sum of the effects of individual systems. Notably, we observed a synergistic protective effect against certain phages even between some mutually exclusive defence systems, such as Zorya II and ietAS. Furthermore, we found that defence systems that are mutually exclusive in one bacterial order can co-occur in others, suggesting that the mutual exclusion between these systems is not caused by mechanistic incompatibility leading to negative epistasis. We also assessed the ecological implications of defence system synergy in short-term competition experiments, finding that bacterial populations carrying synergistic systems gained an evolutionary advantage over populations carrying any of the individual systems when targeted by specific phages that activated the synergistic defence.

Collectively, our results strongly suggest that interactions between defence systems are common and that non-random co-occurrence of defence systems in bacteria is an adaptive phenomenon driven by selection for enhanced immunity against specific phages. The defence strategies appear to differ substantially across bacterial taxa, most likely, driven by the species-specific viromes. Given the extensive horizontal transfer of defence systems, the evolution of bacterial defence strategies appear to follow the venerated general principle of adaptive evolution of microbes “everything is everywhere but the environment selects”^81^.

We further investigated the molecular basis of the synergistic interactions between tmn and its co-occurring defence systems Gabija, Septu I, and PrrC, and uncovered a common mechanism of synergy, whereby tmn co-opted the sensory switch ATPase domains of the companion defence system, enhancing the anti-phage activity. These synergistic interactions and co-opting mechanisms likely play an important role in the evolution of defence systems and the emergence of multiple defence system variants, as observed for CRISPR-Cas^82^, CBASS^19^, Lamassu^6^, Shield^83^, and others. The modularity of anti-phage immunity is further evident in sharing of proteins and functional domains across distinct defence systems^84^, such as HNH-endonuclease in Septu, Zorya II and type II CRISPR-Cas ^2^, NucC in CBASS III^85^ and CRISPR-Cas type III^86^, TOPRIM domain in Gabija^3^, Wadjet^2^, and PARIS^4^, and P-loop NTPases, including helicases, in a broad diversity of defence systems including CRISPR-Cas type I, Gabija, PrrC, tmn and many others^2,3,75^. Expanding our understanding of the mechanisms underlying the modularity of defence systems is expected to provide insights into their adaptive potential and evolutionary dynamics in the perennial arms race between bacteria and phages.

The results of this work underscore the importance of considering the interplay among defence system beyond their cumulative effect, taking into account the environmental context and the influence of phage pressure, for understanding prokaryotic immunity in its increasingly apparent complexity. Adaptation of prokaryotes to specific environments is driven by specific selective forces imposed by the virome, resulting in unique fitness landscapes in each niche. Thus, future research should aim to explore the broader patterns of defence system combinations in diverse prokaryotes in their specific ecological niches.

## Limitations of the study

In this work, we investigated in detail the co-occurrence and mutual exclusion among the known defence systems only among isolates of a single bacterial species, *E. coli*. Our preliminary analysis of defence system co-occurrence in other bacterial orders revealed substantially different patterns emphasizing the importance of a broad exploration of bacterial immunity. Our mechanistic investigation of synergistic defence systems was obviously even more limited so that much further work is required to determine how general the domain co-option observed here might be and what other mechanisms contribute to the synergy.

## ACKNOWLEDGEMENTS

S.K.G and E.V.K. are funded through the Intramural Research Program of the National Institutes of Health of the USA (National Library of Medicine). The work in F.L.N.’s group is supported by Wessex Medical Trust (AB03). We thank Dr Jennifer Mahony (University College Cork) for kindly providing phages JK16 and JK32. We also thank Fagenbank (Netherlands) for providing phages T1, T3, T4, T7, and λ(vir), and Morgen Hedges (University of Southampton) for the phage drawings. We acknowledge the use of the IRIDIS High-Performance Computing Facility, and associated support services at the University of Southampton. We thank Victor Tobiasson for help with Alphafold, and Yuri Wolf and Sanasar Babajanyan for useful discussions.

## AUTHOR CONTRIBUTIONS

Conceptualization, F.L.N.; Methodology, Y.W., S.K.G., Y.E.G., E.V.K., and F.L.N.; Formal Analysis, Y.W., S.K.G., A.v.d.H., and Y.O; Investigation, Y.W., S.K.G., A.v.d.H., C.A.M., S.K., Y.O., C.M.K., and T.T.; Resources, M.C., A.M., Y.E.G., E.V.K., and F.L.N.; Data Curation, Y.W., S.K.G., and F.L.N.; Writing – Original Draft, Y.W. S.K.G., E.V.K., and F.L.N.; Writing – Review & Editing, all authors; Visualization, Y.W., S.K.G., and F.L.N.; Supervision, Y.E.G., E.V.K., and F.L.N.; Funding Acquisition, E.V.K. and F.L.N.

## DECLARATION OF INTERESTS

Y.E.G. is a full-time employee of SNIPR Biome.

## METHODS

### Bacteria and phages

**(A)** *E. coli* strain Dh5α was used to clone plasmids pACYCDuet-1 or 8A with individual defence systems. *E. coli* BL21-AI cells containing plasmid(s) with the defence systems were used for phage assays. All bacterial strains were grown at 37 °C in Lysogeny Broth (LB) with 180 rpm shaking for liquid cultures, or in LB agar (LBA) plates for solid cultures. Strains containing plasmid pACYCDuet-1 or 8A were grown in media supplemented with 25 µg/ml of chloramphenicol or 100 µg/ml of ampicillin, respectively. Phages used in this study and their origins are described in **Table S4**. All phages were produced in LB with their host strain, centrifuged, filter-sterilized, and stored as phage lysates at 4 °C.

### Defence system detection

The FASTA amino acid (FAA), FAST Nucleic Acid (FNA), and Generic Feature Format (GFF) files of 26,384 *Escherichia coli* were downloaded from the NCBI Reference Sequence (RefSeq) Database. Complete genomes of 9,124 Enterobacterales, 1,288 Pseudomonadales, 3,952 Bacillales, and 2,199 Burkholderiales genomes were downloaded from Genbank in October 2021. Defence systems were detected in these genomes using PADLOC version 1.1.0 with database version 1.4.0^87^ and DefenseFinder version 1.0.8^22^. Defence systems outputted by PADLOC in categories *other* and *adaptation* were removed from the analysis. Some of the *E. coli* genomes had extensive fragmentation, which negatively influenced the number of defence systems detected (two-sided Spearman, p < 0.001), but the effect was small enough (r_s_ = -0.06) that we opted to keep the fragmented genomes in our analysis. Defence systems found in all strains can be seen on **Table S1**.

### Contig characterization and prophage detection

Platon version 1.6 ^88^ on accuracy mode was used to categorize bacterial contigs as plasmid or chromosome. Phigaro version 2.2.6^89^ was used with default settings to detect prophages in the bacterial genomes. Contigs shorter than 20 kbp were excluded from this analysis. Defence systems were considered to be located in a prophage region when at least one defence gene was fully within the prophage limits.

### Phylogenetic analysis

Mash v2.3^90^ distances were used to reconstruct phylogenetic trees for each dataset. Pairwise mash distances were calculated, employing a sketch size of 1,000 for the *E. coli* dataset and 100,000 for the order-level datasets. These pairwise distances were transformed into a distance matrix in phylip format, serving as input for the reconstruct of Neighbour-Joining phylogenetic trees with RapidNJ^91^.

To remove potentially contaminated genomes from the phylogenetic trees, we implemented a three-step filtration process. First, we applied TreeShrink v 1.3.9^92^ with centroid rerooting, using a quantile threshold of 0.1 for the *E. coli* dataset and 0.05 for the order-level datasets to remove leaves located on excessively long branches. Second, genomes with CheckM^93^ contamination greater than 5 in the BV-BRC database (https://www.bv-brc.org/) were excluded. Lastly, leaves without metadata in RefSeq or Genbank (as of January 25, 2023), were removed from consideration due to potential errors in their genomic data. After filtration 26,362 genomes of *E. coli* were retained.

For further validation of the phylogenetic trees of the order level datasets, we colour-coded the leaves according to their genus level classification and visually confirmed their agreement with general taxonomy.

To additionally validate the phylogenetic tree for the *E. coli* dataset, we examined the proximity of samples from the same phylogroup the tree. We employed phylogroup assignments from a dataset of 10,667 *E. coli* genomes^45^, and observed that all major clades contained genomes belonging to a single phylogroup. To extract clades containing samples from individual phylogroups, we first employed TreeCluster v 1.0.3^94^ with a threshold parameter of 0.3, resulting in the division of the *E. coli* tree into 17 clusters. Subsequently, we further divided phylogroups E1 and E2, and B1 and C using a custom R script. For each clade, we compiled the list of nodes belonging to that clade and performed phylogroup-specific analysis. All tree visualization and manipulations were performed using R v 4.1.2 with ggtree^95^ and phytools^96^.

### Correlation between defence system content and phylogenetic distance

To investigate the relationship between defence system content and phylogenetic distance we performed a correlation analysis. The phylogenetic distance between pairs of genomes was determined using mash distances obtained as described above. To estimate the distance in defence system content between genomes, we employed Jaccard distance, which compares the presence and absence of defence systems in vectors. Defence systems present in less than 0.5% of genomes in the dataset were excluded from this analysis. For all unique pairs of genomes included in the analysis, we calculated Spearman correlation coefficients, and corresponding p-values, to quantify the correlation between phylogenetic distance and the dissimilarity in defence systems content.

### Odds ratios for defence systems distribution between phylogroups

To test the hypothesis that defence systems are distributed unevenly among *E. coli* phylogroups, we performed a Chi-Squared test for homogeneity. The enrichment analysis aimed to assess whether the presence of a specific defence system in a particular phylogroup deviates from what would be expected by chance. This was quantified by calculating the odds ratio as the ratio of the observed number of genomes containing a particular system within a specific phylogroup to the expected number of genomes with that system.

### Co-occurrence analysis

Given that the genomes under study exhibit phylogenetic relatedness, the co-occurrence analysis required a phylogenetically informed approach. We utilized the Pagel test for binary traits, as implemented in the R-package phytools with fitDiscrete model, to determine if pairs of defence systems exhibited non-random distribution across the phylogenetic tree, indicating potential interdependencies. To ensure robust results and avoid artefacts associated with small sample sizes, systems that were present in less than 0.5% of genomes in the *E. coli* dataset and less than 1% in the order-level datasets were excluded from this analysis. Leaves carrying the defence system of interest were marked with ones, while leaves where the system was absent were marked with zeros. Prior to conducting the test, we standardized tree branch lengths to a mean branch length of 0.1, as recommended in the original implementation of the Pagel test outline in the BayesTraits manual.

We compared the results of the independent model with three alternative models: a) A model in which the distribution of two systems depends on each other; b) A model in which system A depends on system B; c) A model in which system B depends on system A. If model (a) produced a significant p (< 0.01), we further investigated which of the three models best fitted the observed data on the phylogenetic tree. For cases with significant p-value in model (a), we determined the directionality of interaction by using transition probabilities from the best fitted model. This involved two types of transition values: those assuming independent changes of states (e.g. transition from state (0,0) where no system is present, to state (1,0) where system B is present) and those assuming dependent changes of states (e.g. transition from state (0,1) to (1,1)). For each pair of transition values, we calculated the flux, e.g. for the transition from (1,0) to (1,1), by dividing the transition rate from (1,0) to (1,1) by the transition rate from (1,1) to (1,0). If the sum of fluxes into (1,1) exceeded the sum of fluxes from (0,0) to (1,0) and (0,0) to (0,1), we inferred that the systems co-occur; otherwise, they were considered mutually exclusive.

To correct for multiple testing, we used both the Bonferroni correction (the most stringent) and the Benjamini-Hochberg correction (less strict). Both sets of significant results after multiple testing correction were considered for downstream analysis, with a preference for those after Bonferoni correction due to their higher reliability.

### Defence system cloning

The plasmids constructed in this work are listed in **Table S5**, and the primers used can be found in **Table S6**. Plasmid pACYCDuet-1 was modified to contain the pBAD promoter from plasmid 8A (MacroLab) by Gibson assembly. YFP, Druantia III (from *E. coli* ECOR19), ietAS (ECOR52), and tmn (ECOR25) were cloned into the modified pACYCDuet-1 by Gibson assembly. YFP, Kiwa (ECOR8), Gabija (ECOR49), and Zorya II (ECOR19) were cloned into plasmid 8A by Gibson assembly. Coral chromoproteins spisPink and meleRFP (Stanford Free Genes) were cloned into plasmids 8A and pACYC, respectively, by Gibson assembly. The plasmids were recovered in Dh5α cells, extracted using NucleoSpin Plasmid QuickPure Kit (Thermo Fisher Scientific) and confirmed by sequencing at Plasmidsaurus (USA). Mutations of the defence system operons were engineered by around-the-horn PCR, and confirmed by Sanger sequencing (Eurofins Genomics). Plasmids were transformed individually or in combinations into competent BL21-AI cells prepared using the Mix&Go! *E. coli* Transformation Kit (Zymo).

### Efficiency of platting

Overnight cultures of the bacteria were diluted 1:50 in LB containing antibiotics, induced with 0.2% arabinose, and incubated for 5 hours before being used in double agar overlay assays. For this, bacterial cultures were mixed with 0.6% top agar and overlaid on LBA plates. Ten-fold serial dilutions of the phage stocks were spotted onto the bacterial lawn and the plates incubated overnight at 37 °C. The phage plaques were counted and used to calculate the EOP relative to the control. Epistatic coefficients, representing the interaction strength between defence systems, were determined as |Log10 (EOP_system1+system2_)| – |Log10 (EOP_system1_)| – |Log10 (EOP_system2_)|. Synergy was considered when |Log10 (EOP_system1+system2_)| > |Log10 (EOP_system1_)| + |Log10 (EOP_system2_)| +1, additive effects were considered when Max [|Log10 (EOP_system1_)|, |Log10 (EOP_system2_)|] < |Log10 (EOP_system1+system2_)| < |Log10 (EOP_system1_)| + |Log10 (EOP_system2_)| + 1, and antagonistic effects were considered when |Log10 (EOP_system1+system2_)| – Max (|Log10 (EOP_system1_)|, |Log10 (EOP_system2_)|] < -1. Statistical significance was determined using the multiple comparison function from Two-way ANOVA with a p-value of <0.01.

### Time post infection assay

Overnight bacterial cultures were diluted to an optical density at 600 nm of 0.1 in LB containing antibiotics and 0.2% arabinose. The cultures were infected with phage at an MOI of 0.0001. At 0, 1, 2, 3 and 4 hours post infection, a sample was taken and centrifuged at 12,000 × *g* for 2 minutes. The supernatant was serially diluted, and the phages were quantified by plaque assay on a bacterial lawn of cells with YFP. PFUs were counted after overnight incubation at 37 °C.

### Liquid assay

Overnight bacterial cultures were diluted to an optical density at 584 nm of 0.25 in LB containing antibiotics and 0.2% arabinose. The bacterial suspension was distributed into wells of a 96-well plate to which phage dilutions or LB were added. The plates were incubated in a Fluostar Optima plate reader at 37 °C with shaking at 200 rpm, with optical density at 584 nm measured every 5 min for 24 h. To evaluate the impact of interactions between defence systems, we calculated the areas under the curve (AUC) for both individual systems and combination of systems. To calculate the AUC, the optical density at the start of the experiment was subtracted from each data point. If the AUC for a system combination exceeded the sum of the AUCs for the individual systems (the expected value), we considered those system as having a synergistic protective effect.

### Short-term evolution experiment

Overnight bacterial cultures with single or double defence systems were diluted to an optical density at 600 nm of 0.1 in LB containing antibiotics and 0.2% arabinose, and mixed in equal proportions. The mixed cultures were infected with phage at an MOI of 0.0001, and incubated at 37 °C with shaking at 180 rpm for 24 h. A control without phage was used. The cultures were centrifuged at 8000 × *g* for 10 min, and the cell pellet was washed three times with LB at 12,000 × *g* for 2 min. Cells were resuspended in 1 ml of LB, serially diluted, and 100 µl of each dilution were spread onto LBA plates supplemented with antibiotics and 0.2% L-arabinose. The plates were incubated overnight at 37 °C and the colonies of each colour counted. The experiment was repeated for 48h and 72h time points, using 50 µl of the previous cultures to inoculate fresh LB containing antibiotics and 0.2% arabinose, and challenging the cultures with phage at the same MOI.

### Quantification of receptor mutants

Ten colonies were selected for each time point and condition of the short-term evolution experiment. The selected colonies were resuspended in 30 µl of sterile ddH_2_O, and 5 µl of this cell suspension were spotted onto LBA plates. To assess the presence of receptor mutants, 2 µl of phage stock were spotted on top of the bacteria spots. The plates were left to incubate overnight at 37 °C. Colonies where no evidence of phage lysis was observed were identified and counted as receptor mutants.

### Plasmid loss assay

Overnight bacterial cultures containing double defence systems were diluted to an optical density at 600 nm of 0.1 in LB containing 0.2% arabinose. The cultures were infected with phage at an MOI of 0.0001 and incubated at 37 °C with shaking at 180 rpm for 24h. A control without phage was used. Cultures were centrifuged at 8000 × *g* for 10 min, and the cell pellet was washed three times with LB at 12,000 × *g* for 2 min. Cells were resuspended in 1 ml of LB, serially diluted, and 100 µl of each dilution were spread onto LBA plates. The plates were incubated overnight at 37 °C and 96 colonies were picked, dissolved in LB, and streaked onto LBA plates with and without antibiotics, to determine the rate of plasmid loss. The experiment was repeated for 48h and 72h time points, using 50 µl of the previous cultures to inoculate fresh LB containing 0.2% arabinose, and challenging the cultures with phage at the same MOI.

### Cactus plasmid analysis

*E. coli* plasmids containing both tmn and Gabija defence systems were extracted from the Enterobacterales dataset. The plasmid dataset comprised of a total of 104 plasmids from various *E. coli* strains. Next, we constructed a phylogenetic tree for these plasmids using the Neighbour-Joining method as described above, using a mash sketch size of 1,000. Next, we generated whole length plasmid alignments using Cactus v 2.4.4^97^ with default parameters and the phylogenetic tree described above as the guiding tree. The reference free alignment obtained was then transformed into multiple alignment format (MAF) using plasmid CP083423.1 as a reference. This MAF file was filtered and visualized in R. Alignment blocks smaller than 100 bp were excluded from the analysis. Alignment blocks were considered to be present in the plasmid if they exhibited a coverage above 50%.

## DATA AVAILABILITY

All unique bacterial strains, phages, and plasmids generated in this study are available from the lead contact without restriction. Phages of SNIPR Biome are proprietary and can be shared with other non-competing parties upon written permission. Raw data have been deposited at Github (https://github.com/garushyants/synergy_bacterial_immune_systems/) and Zenodo (https://doi.org/10.5281/zenodo.10075784) and are publicly available as of the date of publication.

## QUANTIFICATION AND STATISTICAL ANALYSIS

A two-sided binomial test was performed to determine if the observed co-occurrence of defence systems differed significantly from the expected co-occurrence, using R^98^. To correct for multiple testing we utilized both Benjamin-Hochberg and Bonferroni corrections, with a p-value < 0.001 considered significant. Unless stated otherwise, experimental data are presented as the mean of biological triplicates ± standard deviation. Statistical tests were performed using GraphPad Prism 9.2.0 and one sample t test or one-way ANOVA test. All statistical tests are described in detail in the corresponding chapters in Methods, and available as R code on Github and Zenodo.

## SUPPLEMENTAL INFORMATION TITLES AND LEGENDS

**Figure S1** Distribution of defence systems across *Escherichia coli* genomes.

**(A)** Phylogenetic tree of the *E. coli* genomes, with the count of defence systems per genome displayed and colour-coded based on their genomic locations. Each phylogroup is colour-coded according to the key.
**(B)** Distribution pattern of defence systems within the chromosome, plasmids, and prophage regions of *E. coli*.
**(C)** Density plot demonstrating the correlation between mash distance and similarity in defence systems content among the *E. coli* strains. A linear regression trend line is highlighted in red for reference.

**Figure S2** Counts, genomic distance, and location of co-occurring defence systems in *E. coli*.

**(A)** Network of co-occurrence between defence systems. The width of the edge is proportional to the number of genomes in which the pair was observed, and the size of the node is proportional to the number of genomes where the specific defence system was found. Co-occurrences are shown in orange, while mutual exclusions are shown in green.
**(B)** Counts of co-occurrences and mutual exclusions for each defence system. Correlation between pairs was assessed using the Pagel test for binary traits. Significant correlations, adjusted with either the Benjamini-Hochberg or the Bonferroni method, are indicated by the colour scheme in the legend.
**(C)** Genomic distance between pairs of co-occurring defence systems.
**(D)** Distribution of co-occurring pairs across chromosome, plasmid, or prophage regions. Instances where each defence system resides in different locations are denoted as Plasmid-Chromosome, Prophage-Chromosome, and Prophage-Plasmid.

**Figure S3** Synergistic anti-phage defence provided by combinations of defence systems.

**(A)** Efficiency of plating (EOP) of phages on strains carrying individual defence systems and their combinations. b.d., below limit of detection. Unfilled circles indicate instances where it was not possible to determine the number of phage plaques, hence a value of 1 at the respective dilution was assumed. Asterisks (*, p< 0.01; **, p< 0.001) indicate cases of synergy.
**(B)** Time post infection assays, measuring phage tires over the course of four hours in liquid cultures of *E. coli* containing individual or combined defence systems. Data is shown as the average and standard deviation of three biological replicates.
**(C)** Bacterial growth under phage predation at different multiplicities of infection (MOIs), represented as area under the curve (AUC) in OD⋅h. A defence system pair acts synergetic when its dot (red) is above the expected additive effect (blue). Data is shown as the confidence interval of three biological replicates. The raw data and growth curves used to calculate the AUCs are available on the associated Github and Zenodo databases.

**Figure S4** Comparative analysis of defence systems across bacterial orders.

**(A)** Prevalence of defence systems in the bacterial orders Pseudomonadales, Enterobacterales, Burkholderiales, and Bacillales. The defence systems are colour-coded according to their mechanism of defence.
**(B)** Venn diagram illustrating the overlap in defence system presence among the four bacterial orders and *E. coli*.

**Figure S5** Assessment of dual plasmid maintenance in *E. coli* using plasmid loss assays.

**(A)** Illustration outlining the plasmid loss assay procedure. Bacterial cultures harbouring two plasmids with one defence system each, were grown without antibiotic and assessed for plasmid persistence every 24h for three days. Plasmid maintenance was determined by spreading serial dilutions of the bacterial cultures onto LBA plates. Subsequently, 100 colonies were streaked on LBA with and without corresponding antibiotics. This assay was performed both in the absence and presence of phage.
**(B)** Proportion of colonies maintaining either one or both plasmids in the plasmid loss assay under normal conditions (control) and under phage predation.

**Figure S6** Insights into tmn and its synergistic interaction with Gabija and Septu I.

**(A)** Tertiary structure of the tmn protein predicted by Alphafold2 (pLDDT 89.3). The figure depicts two chains of the potential hexamer complex. Light purple, P-loop NTPase; dark grey/beige, predicted transmembrane regions; pink, Walker A motif; purple, Walker B motif;

The P-loop NTPase domain is shown in light purple, the predicted transmembrane regions is shown in dark grey/beige. Residues mutated in the experiments are shown as spheres.

**(B)** Efficiency of plating (EOP) of phages T1, 670, and 678 on cells expressing tmn (T), Septu I (S), Gabija (G), Septu and tmn (ST), or Gabija and tmn (GT), derived from different strains. Unfilled circles represent cases where the number of phage plaques was indeterminable, with a value of 1 assumed at the respective dilution. Asterisks (**) denote cases of synergy.

**Table S1** Defence systems found per strain of *E. coli*, Bacillales, Burkholderiales, Enterobacterales, and Pseudomonadales.

**Table S2** Enrichment analysis of defence systems in *Escherichia coli* phylogroups, using the chi-squared test for homogeneity. Only systems exhibiting significant enrichment (p-value <0.001) are shown

**Table S3** Co-occurrence analysis of defence systems in *E. coli*, Bacillales, Burkholderiales, Enterobacterales, and Pseudomonadales.

**Table S4** List of phages used in this work.

**Table S5** List of plasmids used and generated in this work.

**Table S6** List of primers used in this work.

## Notes

https://github.com/garushyants/synergy_bacterial_immune_systems/

https://doi.org/10.5281/zenodo.10075784

### Summary of Updates

Data analysis exploring the correlations between the occurrences of pairs of defence systems in E. coli genomes corrected for phylogenetic bias. Additional experimental data to validate and explain the synergy between defence systems.

