## Supplemental information for "Synergistic anti-phage activity of bacterial defence systems"

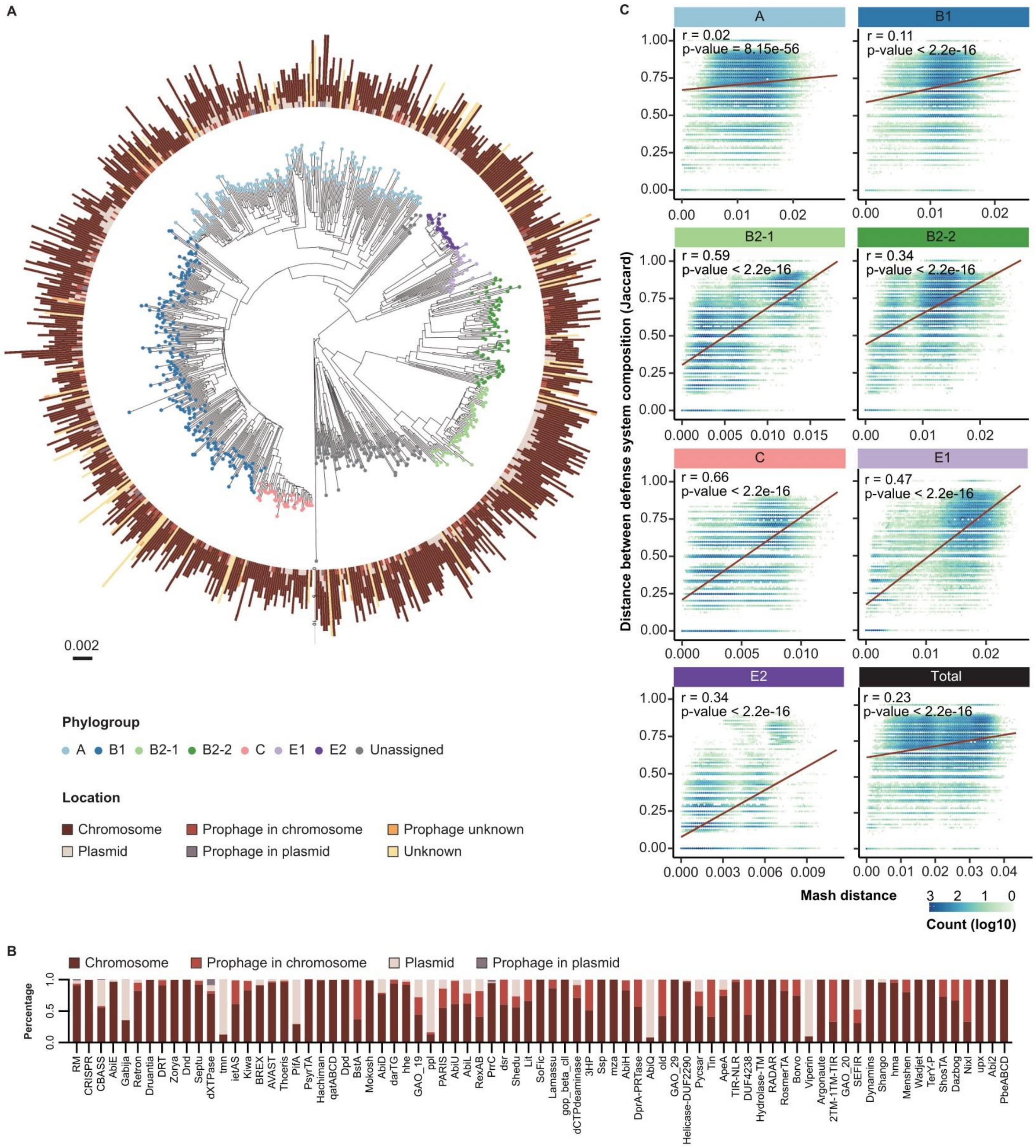


**Figure S1** Distribution of defence systems across *Escherichia coli* genomes.

**(A)** Phylogenetic tree of the *E. coli* genomes, with the count of defence systems per genome displayed and colour-coded based on their genomic locations. Each phylogroup is colour-coded according to the key.

**(B)** Distribution pattern of defence systems within the chromosome, plasmids, and prophage regions of *E. coli*.

**(C)** Density plot demonstrating the correlation between mash distance and similarity in defence systems content among the *E. coli* strains. A linear regression trend line is highlighted in red for reference.

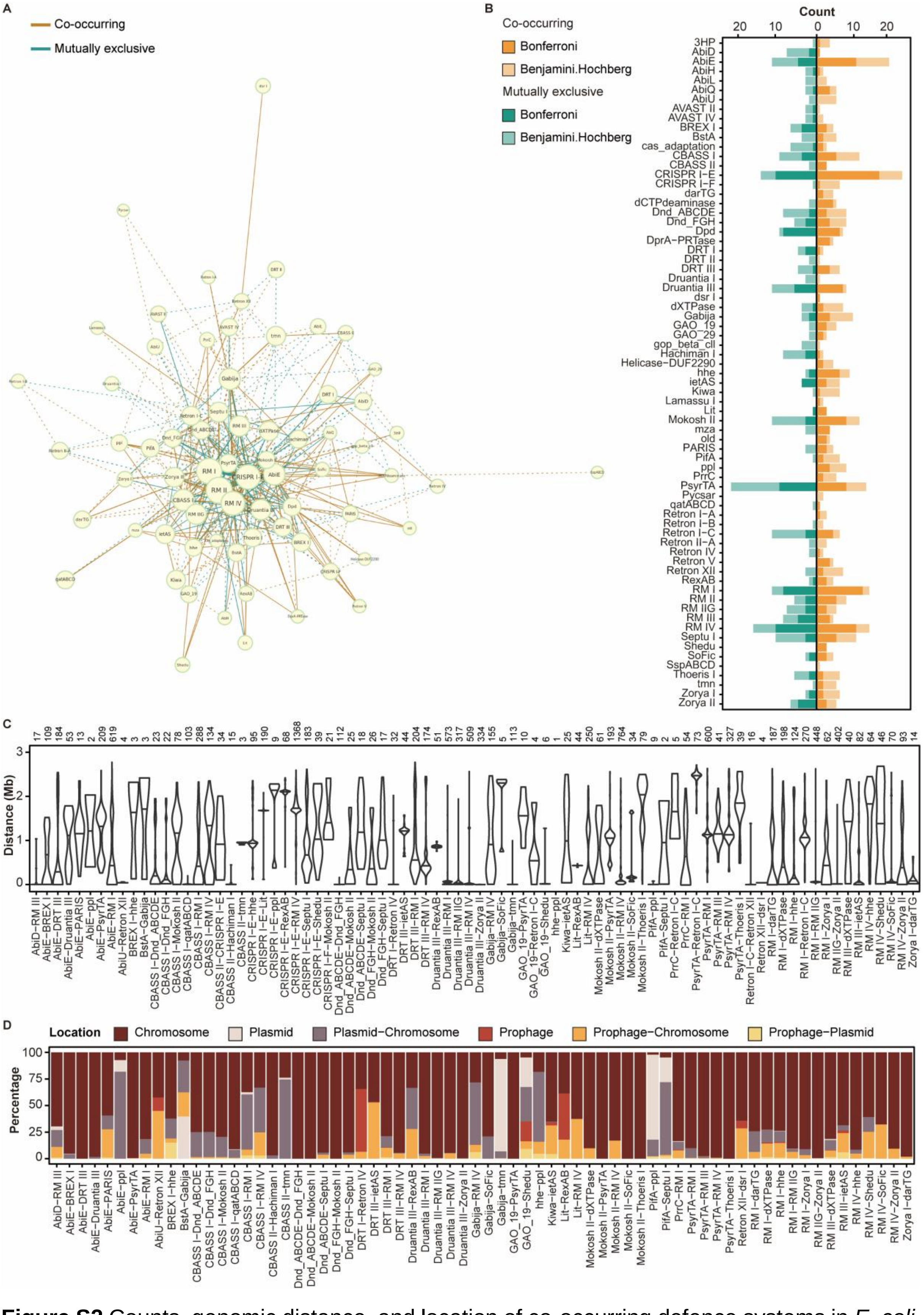


**Figure S2** Counts, genomic distance, and location of co-occurring defence systems in *E. coli*. **(A)** Network of co-occurrence between defence systems. The width of the edge is proportional to the number of genomes in which the pair was observed, and the size of the node is proportional to the number of genomes where the specific defence system was found. Co-occurrences are shown in orange, while mutual exclusions are shown in green.

**(B)** Counts of co-occurrences and mutual exclusions for each defence system. Correlation between pairs was assessed using the Pagel test for binary traits. Significant correlations, adjusted with either the Benjamini-Hochberg or the Bonferroni method, are indicated by the colour scheme in the legend.

**(C)** Genomic distance between pairs of co-occurring defence systems.

**(D)** Distribution of co-occurring pairs across chromosome, plasmid, or prophage regions. Instances where each defence system resides in different locations are denoted as Plasmid-Chromosome, Prophage-Chromosome, and Prophage-Plasmid.

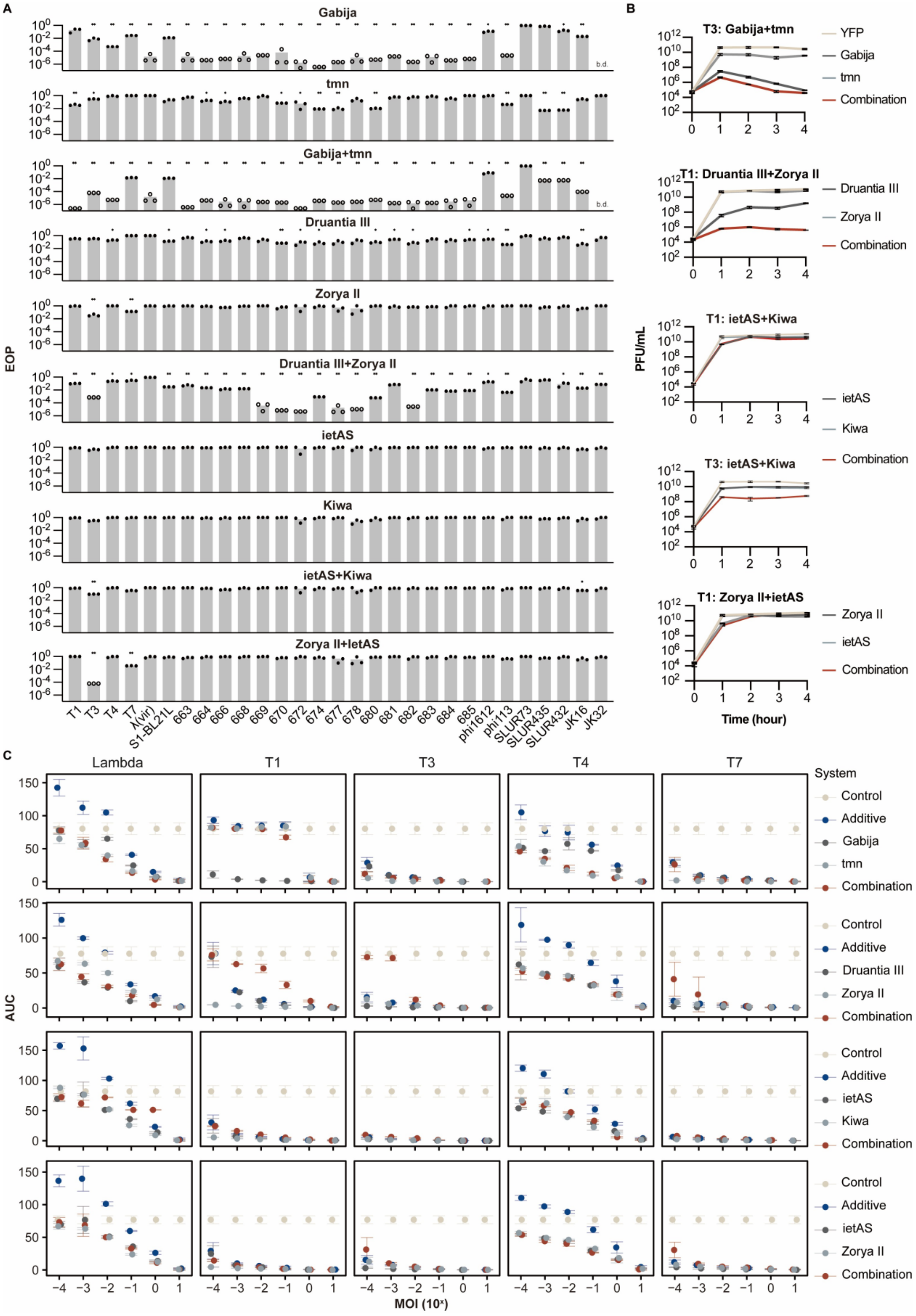


**Figure S3** Synergistic anti-phage defence provided by combinations of defence systems.

**(A)** Efficiency of plating (EOP) of phages on strains carrying individual defence systems and their combinations. b.d., below limit of detection. Unfilled circles indicate instances where it was not possible to determine the number of phage plaques, hence a value of 1 at the respective dilution was assumed. Asterisks (*, p< 0.01; **, p< 0.001) indicate cases of synergy.
**(B)** Time post infection assays, measuring phage tires over the course of four hours in liquid cultures of *E. coli* containing individual or combined defence systems. Data is shown as the average and standard deviation of three biological replicates.
**(C)** Bacterial growth under phage predation at different multiplicities of infection (MOIs), represented as area under the curve (AUC) in OD·h. A defence system pair acts synergetic when its dot (red) is above the expected additive effect (blue). Data is shown as the confidence interval of three biological replicates. The raw data and growth curves used to calculate the AUCs are available on the associated Github and Zenodo databases.

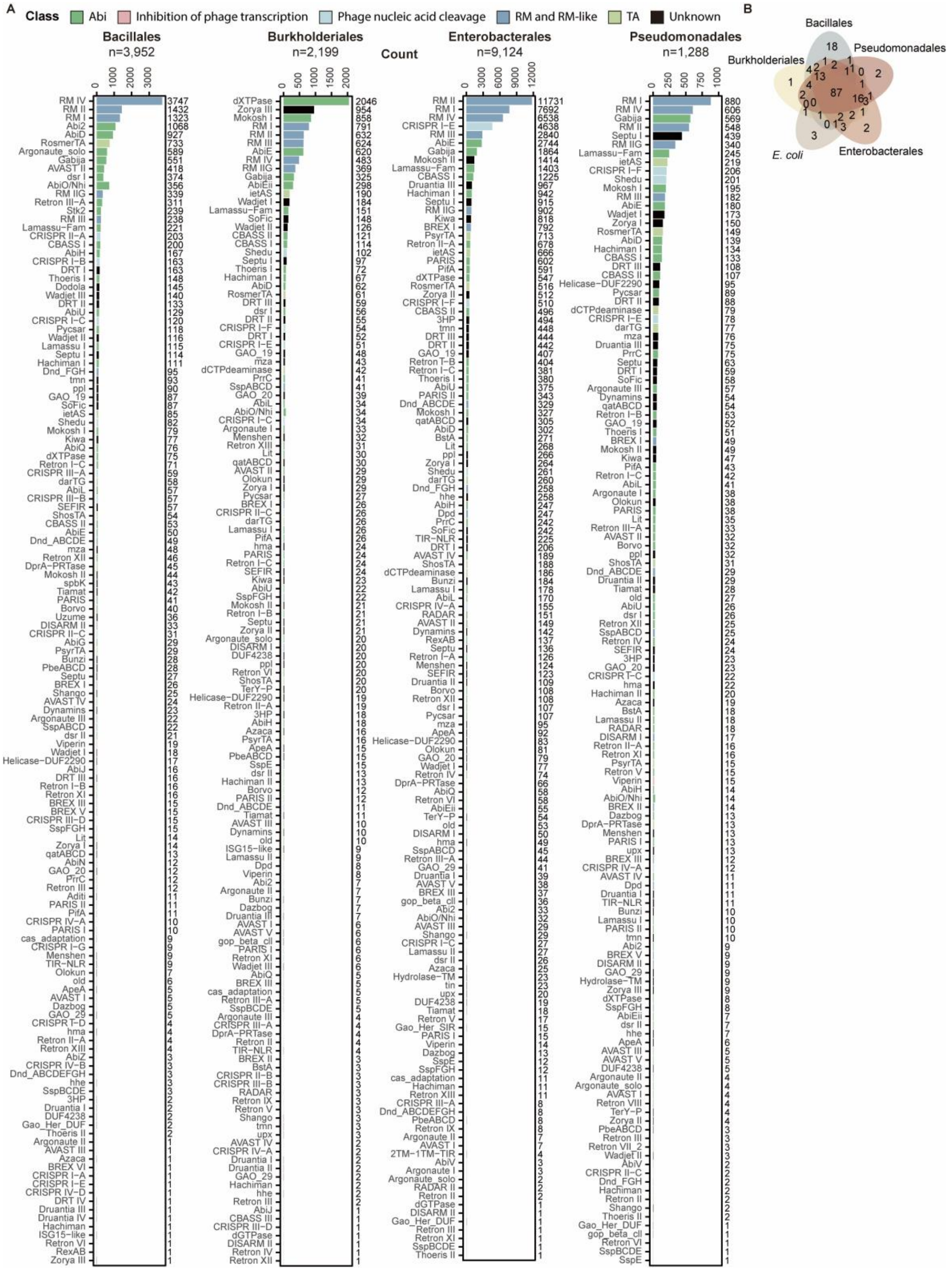


**Figure S4** Comparative analysis of defence systems across bacterial orders.

**(A)** Prevalence of defence systems in the bacterial orders Pseudomonadales, Enterobacterales, Burkholderiales, and Bacillales. The defence systems are colour-coded according to their mechanism of defence.

**(B)** Venn diagram illustrating the overlap in defence system presence among the four bacterial orders and *E. coli*.

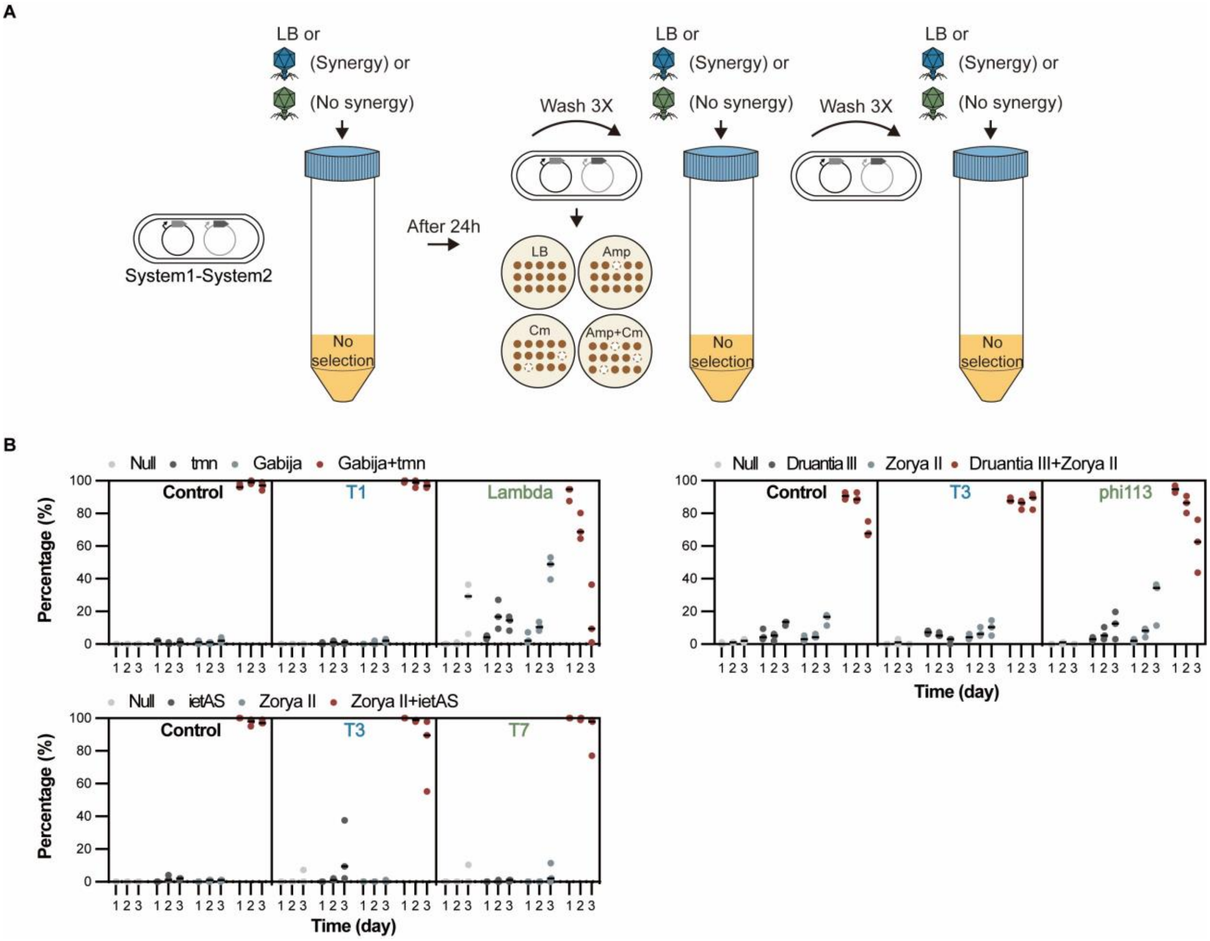


**Figure S5** Assessment of dual plasmid maintenance in *E. coli* using plasmid loss assays.

**(A)** Illustration outlining the plasmid loss assay procedure. Bacterial cultures harbouring two plasmids with one defence system each, were grown without antibiotic and assessed for plasmid persistence every 24h for three days. Plasmid maintenance was determined by spreading serial dilutions of the bacterial cultures onto LBA plates. Subsequently, 100 colonies were streaked on LBA with and without corresponding antibiotics. This assay was performed both in the absence and presence of phage.

**(B)** Proportion of colonies maintaining either one or both plasmids in the plasmid loss assay under normal conditions (control) and under phage predation.

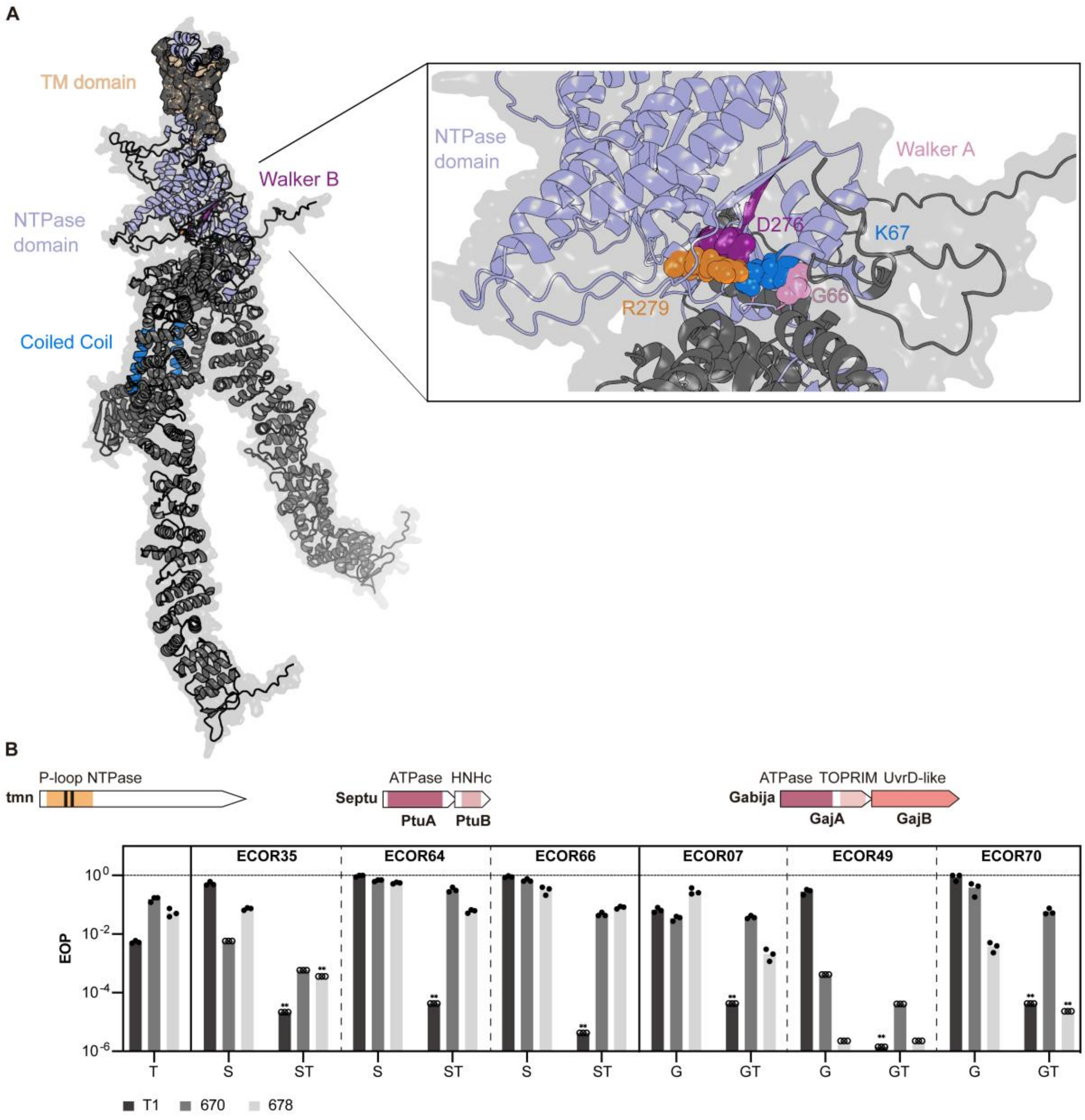


**Figure S6** Insights into tmn and its synergistic interaction with Gabija and Septu I.

**(A)** Tertiary structure of the tmn protein predicted by Alphafold2 (pLDDT 89.3). The figure depicts two chains of the potential hexamer complex. Light purple, P-loop NTPase; dark grey/beige, predicted transmembrane regions; pink, Walker A motif; purple, Walker B motif; The P-loop NTPase domain is shown in light purple, the predicted transmembrane regions is shown in dark grey/beige. Residues mutated in the experiments are shown as spheres.

**(B)** Efficiency of plating (EOP) of phages T1, 670, and 678 on cells expressing tmn (T), Septu I (S), Gabija (G), Septu and tmn (ST), or Gabija and tmn (GT), derived from different strains. Unfilled circles represent cases where the number of phage plaques was indeterminable, with a value of 1 assumed at the respective dilution. Asterisks (**) denote cases of synergy.

**Table S2** Enrichment analysis of defence systems in *Escherichia coli* phylogroups, using the chi-squared test for homogeneity. Only systems exhibiting significant enrichment (p-value <0.001) are shown

| Defence system | Chi-squared | p.value | Odds ratio on phylogroup | | | | | | |
|---|---|---|---|---|---|---|---|---|---|
| | | | A | B1 | B2.1 | B2.2 | C | E1 | E2 |
| Zorya II | 14969.9 | 0.0005 | 0.4585 | 0.1269 | 0.0000 | 0.0465 | 0.0095 | 4.6239 | 10.2519 |
| Druantia III | 9333.2 | 0.0005 | 0.7045 | 0.3891 | 0.0168 | 0.2745 | 0.0077 | 4.8652 | 6.9160 |
| Retron I-C | 13994.9 | 0.0005 | 0.1854 | 0.3406 | 9.0435 | 0.0998 | 0.0204 | 0.1162 | 0.0000 |
| AbiE | 8028.8 | 0.0005 | 0.1656 | 0.9016 | 4.7719 | 0.3916 | 1.8202 | 1.8051 | 0.0083 |
| CRISPR I-F | 1893.8 | 0.0005 | 0.0000 | 0.9783 | 0.0000 | 5.1161 | 0.0000 | 0.0346 | 0.0000 |
| Thoeris I | 5973.0 | 0.0005 | 0.3762 | 0.1756 | 0.2708 | 6.1609 | 0.0438 | 0.0332 | 0.0000 |
| Septu I | 1703.8 | 0.0005 | 0.4943 | 0.9695 | 0.1499 | 3.1719 | 0.0955 | 2.2554 | 0.0000 |
| qatABCD | 2495.1 | 0.0005 | 0.7275 | 0.4094 | 0.2379 | 4.6304 | 0.0349 | 0.4959 | 0.0425 |
| BREX I | 784.3 | 0.0005 | 0.6787 | 1.2150 | 1.3696 | 0.3668 | 3.5390 | 1.1264 | 0.0000 |
| PsyrTA | 5636.9 | 0.0005 | 0.0156 | 0.0333 | 3.2218 | 1.6658 | 0.0000 | 0.1392 | 7.8123 |
| RM IV | 1398.5 | 0.0005 | 1.2216 | 0.8888 | 0.7879 | 1.6031 | 0.4293 | 1.0178 | 0.0242 |
| RM IIG | 10256.3 | 0.0005 | 0.8956 | 0.4458 | 0.1013 | 0.2037 | 0.3210 | 1.4935 | 7.8641 |

**Table S4** List of phages used in this work

| Phage | Morphology | Accession no. | Provided by |
|---|---|---|---|
| T1 | Siphophage | NC_005833.1 | Fagenbank |
| T3 | Podophage | NC_003298.1 | Fagenbank |
| T4 | Myophage | NC_000866.4 | Fagenbank |
| T7 | Podophage | NC_001604.1 | Fagenbank |
| λ(vir) | Siphophage | NC_001416 [a] | Fagenbank |
| S1-BL21L | Myophage | NS | Fagenbank |
| T7Select | Podophage | [b] | Novogen |
| phi1612 | Podophage | NS | Martha Clokie |
| phi113 | Podophage | NS | Martha Clokie |
| SLUR435 | Podophage | NS | Andrew Millard |
| SLUR432 | Podophage | NS | Andrew Millard |
| JK16 | Siphophage | MK962751 | Jennifer Mahony |
| JK32 | Myophage | MK962753 | Jennifer Mahony |
| 663 | Myophage | NS | SNIPRBiome |
| 664 | Myophage | NS | SNIPRBiome |
| 666 | Myophage | NS | SNIPRBiome |
| 668 | Myophage | NS | SNIPRBiome |
| 670 | Myophage | NS | SNIPRBiome |
| 672 | Myophage | NS | SNIPRBiome |
| 674 | Myophage | NS | SNIPRBiome |
| 677 | Myophage | NS | SNIPRBiome |
| 678 | Myophage | NS | SNIPRBiome |
| 680 | Myophage | NS | SNIPRBiome |
| 681 | Myophage | NS | SNIPRBiome |
| 682 | Myophage | NS | SNIPRBiome |
| 683 | Myophage | NS | SNIPRBiome |
| 684 | Myophage | NS | SNIPRBiome |
| 685 | Myophage | NS | SNIPRBiome |

NS, not sequenced

[a] Mutation in *cI* [1]

[b] can be downloaded from https://www.merckmillipore.com/INTL/en/product/T7Select-415-1-DNA,EMD_BIO-70040?ReferrerURL=https%3A%2F%2Fwww.google.com%2F#anchor_VSEQ

**Table S5** List of plasmids used and generated in this work

| Name | Code | Insert | Derived from | Resistance |
|---|---|---|---|---|
| pACYCDuet-1 | pUOS001 | - | - | $Cm^R$ |
| 8A | pUOS041 | - | - | $Amp^R$ |
| pCOLA | pUOS042 | - | - | $Kan^R$ |
| pCDF | pUOS084 | - | - | $Spc^R$ |
| pACYC | pUOS043 | pBAD promoter from 8A | pACYCDuet-1 | $Cm^R$ |
| pACYC-YFP | pUOS031 | YFP | pACYC | $Cm^R$ |
| pIetAS | pUOS032 | ietAS from ECOR52 | pACYC | $Cm^R$ |
| pKiwa | pUOS033 | Kiwa from ECOR8 | 8A | $Amp^R$ |
| pDruantiaIII | pUOS034 | Druantia III from ECOR19 | pACYC | $Cm^R$ |
| pZoryaII | pUOS035 | Zorya II from ECOR19 | 8A | $Amp^R$ |
| pTmn | pUOS036 | tmn from ECOR25 | pACYC | $Cm^R$ |
| pGabija | pUOS037 | Gabija from ECOR49 | 8A | $Amp^R$ |
| 8A-YFP | pUOS038 | YFP | 8A | $Amp^R$ |
| pCOLA-YFP | pUOS039 | YFP | pCOLA | $Kan^R$ |
| pGabija GajA E379A | pUOS063 | GajA E379A | pGabija | $Amp^R$ |
| pGabija GajA D495A | pUOS064 | GajA D495A | pGabija | $Amp^R$ |
| pGabija GajA K528A | pUOS065 | GajA K528A | pGabija | $Amp^R$ |
| pGabija GajA H317A | pUOS066 | GajA H317A | pGabija | $Amp^R$ |
| pGabija GajA K35A | pUOS067 | GajA K35A | pGabija | $Amp^R$ |
| pGabija GajB D183A/E184A | pUOS068 | GajB D183A/E184A | pGabija | $Amp^R$ |
| pGabija GajB K27A/T28A | pUOS069 | GajB K27A/T28A | pGabija | $Amp^R$ |
| pTmn G66A/K67A | pUOS076 | tmn G66A/K67A | pTmn | $Cm^R$ |
| pTmn D276A/R279A | pUOS078 | tmn D276A/R279A | pTmn | $Cm^R$ |
| 8A-spisPink | pUOS082 | - | 8A | $Amp^R$ |

| | | | | |
|---|---|---|---|---|
| pACYC-meleRFP | pUOS083 | - | pACYC | $Cm^R$ |
| pGajA | pUOS103 | Stop codon at amino acid 13-15 on GajB | pGabija | $Amp^R$ |
| pGajB | pUOS104 | Stop codon at amino acid 346, 350 and 353 on GajA | pGabija | $Amp^R$ |
| pSeptuI | pUOS086 | Septu I from ECOR35 | 8A | $Amp^R$ |
| pSeptuI PtuA K43A | pUOS095 | PtuA K43A | pSeptuI | $Amp^R$ |
| pSeptuI PtuA D316A/E317A | pUOS097 | PtuA D316A/E317A | pSeptuI | $Amp^R$ |
| pSeptuI PtuA H321A | pUOS098 | PtuA H321A | pSeptuI | $Amp^R$ |
| pSeptuI PtuB K6A | pUOS100 | PtuB K6A | pSeptuI | $Amp^R$ |
| pSeptuI PtuB H75A | pUOS101 | PtuB H75A | pSeptuI | $Amp^R$ |
| pPrrC | pUOS102 | PrrC from bSNP1527 | 8A | $Amp^R$ |

**Table S6** List of primers used in this work

| Primer | Sequence (5' – 3') | Description |
|---|---|---|
| *Plasmid backbone* | | |
| FN0407 | ATCCCAACTCCATAAGGATC | Amplification of pBAD LIC-8A backbone |
| FN0408 | ATCTATATCTCCTTCTTAAAGTTAAACA | |
| FN0409 | CGACTCCTGCATTAGGAAATAAGAAACCAATTGTCCATATTGC | Amplification of pBAD promoter (with FN0408) |
| FN0410 | TCTACTAGCGCAGCTTAATTAAC | Amplification of pACYCDuet-1 backbone |
| FN0411 | ATTTCCTAATGCAGGAGTCGC | |
| FN0418 | TTTAAGAAGGAGATATAGATATGGTGAGCAAGGGCGAG | Amplification of YFP from pACYCDuet-1-YFP |
| FN0419 | TTATGGAGTTGGGATTTACTTGTACAGCTCGTCCATGC | |
| FN0484 | GCCTCTAAACGGGTCTTGAG | Amplification of pCOLA backbone |
| FN0485 | ATTTCGATTATGCGGCCGTGTA | |
| FN0841 | GCCTCTAAACGGGTCTTGTTAACCTAGGCTGCTGCCA | Amplification of pCDF backbone |
| FN0842 | TTGGTAACGAATCAGACAATGCGCAACGCAATTAATGTAAGTTAG | |
| *Defence systems* | | |
| FN0376 | TTTAAGAAGGAGATATAGATCTGCCTTCCTTTGATACAATT | Amplification of Zorya II from ECOR19 |
| FN0366 | TTATGGAGTTGGGATCTTATTATATTGCTTTATTCATTGTGACTGTC | |
| FN0377 | TTTAAGAAGGAGATATAGATCTCGAGCTGGTGAGTCGTTA | Amplification of Kiwa from ECOR8 |
| FN0398 | TTATGGAGTTGGGATCTTATTATACTGAACAATATTATCTGCATCCCTTG | |
| FN0399 | TTTAAGAAGGAGATATAGATCATGAGAATTGAAACTGTTTATATTAAAGGATTC | Amplification of Gabija from ECOR49 |
| FN0400 | TTATGGAGTTGGGATCTTATTAAACTCCCCGAAAAATCCG | |
| FN0412 | TTTAAGAAGGAGATATAGATTTTGATACCTGTGTAAATAATGGA | Amplification of Druantia III from ECOR19 |
| FN0413 | ATTAAGCTGCGCTAGTAGACGGTTTTTCTGTCTATTTGGG | |
| FN0414 | AAGAAGGAGATATAGATATAGAACGATGAAGGATGGAAGC | Amplification of ietAS from ECOR52 |
| FN0415 | AATTAAGCTGCGCTAGTAGATTCTTAATCTCTCGATGGCGC | |
| FN0416 | TTTAAGAAGGAGATATAGATAAAATCCCCACGGCAGGC | Amplification of tmn from ECOR25 |
| FN0417 | AATTAAGCTGCGCTAGTAGATTTCCGGAAGTGTACCACGC | |
| FN0713 | TTTAAGAAGGAGATATAGATCATCATGACGGGCATACGCTTTC | Amplification of Septu I from ECOR35 |
| FN0714 | TTATGGAGTTGGGATCGCATACACCACACGACG | |
| FN8516 | TTTAAGAAGGAGATATAGATCCTGACTCCATCACCGAAG | Amplification of PrrC from bSNP1527 |
| FN8517 | ATCCCAACTCCATAATCAACCATTTTTTTGTTCCTGCT | |
| *Insert* | | |
| FN0483 | CACGGCCGCATAATCGAAATAAGAAACCAATTGTCCATATTGC | Amplification of YFP from pACYC-YFP |
| FN0490 | CAAGACCCGTTTAGAGGCTTACTTGTACAGCTCGTCCAT | |
| FN0520 | TTTAAGAAGGAGATATAGATATGTCGCACTCAAAACAAGCA | Amplification of spiPink |

| Primer | Sequence (5' – 3') | Description |
|---|---|---|
| FN0808 | CCTTATGGAGTTGGGATTCACACTTCCAGCACACG | |
| FN0797 | TTTAAGAAGGAGATATAGATATGAGCGTTATTAAGCCGGA | Amplification of meleRFP |
| FN0798 | AATTAAGCTGCGCTAGTAGACGGTCTCAAAGCTCTTCACTG | |
| *Defence system mutants* | | |
| FN0623 | P-TTTTTTGGTCGCGGGACCATCTG | Amplification of pGabija for mutation GajA E379A |
| FN0624 | P-ACCACGTTAGCAAAAAATG | |
| FN0625 | P-ATCTTTTTTTGCTCTTGAAGGAGATCTTG | Amplification of pGabija for mutation GajA D495A |
| FN0626 | P-ATATAAACGCCGTTAGGATTTATTATTG | |
| FN0627 | P-GAGAGGAAAAGCAGCTATACGTATGC | Amplification of pGabija for mutation GajA K528A |
| FN0628 | P-AAATAATCAGCGGCATCATC | |
| FN0629 | P-CGTTACAAGCGCCTCACCTCAAATTGC | Amplification of pGabija for mutation GajA H317A |
| FN0630 | P-ATTGCTTGGCCTGGAAGT | |
| FN0631 | P-TGATGTTGGAGCGACCAACCTATTATATG | Amplification of pGabija for mutation GajA K35A |
| FN0632 | P-TTTGCACCTATAATCAAATTAG | |
| FN0633 | P-TTTTTTTGTCGCTGCGTATCAGGATAC | Amplification of pGabija for mutation GajB D183A/E184A |
| FN0634 | P-CCAATATACTTTGCCATTAAATAC | |
| FN0635 | P-AGGAAGTGGTGCAGCAACAATACTCG | Amplification of pGabija for mutation GajB K27A/T28A |
| FN0636 | P-GGGCATGCAGTTATAACC | |
| FN0684 | P-GTATGGGGCAGCGGCAAGCTCAGTATTAAAAAC | Amplification of pTmn for mutation G66A/K67A |
| FN0685 | P-GGCCCGGTAACAGCAATA | |
| FN0688 | P-GACGCTTTTAACAATGGCCGGATTTTC | Amplification of pTmn for mutation D276A/R279A |
| FN0689 | P-AAGAGCTTCGAATATCACTACATCATATTTAG | |
| FN0835 | P-TAGTAATGAGGTAATCTGGTTATAACTGCATG | Amplification of pGabija for introducing stop codon in GajB |
| FN0836 | P-ATCAATTGCATCTAACTGTTCCTTAG | |
| FN0837 | P-TAATGATAAGCCGCAAGTAATGGATGTTC | Amplification of pGabija for introducing stop codon in GajA |
| FN0838 | P-TAGAGTCTCTCCATCATCATAGT | |
| FN0899 | ATTAAAATCGTTGGCAACCAGC | Amplification of pSeptuI for mutation PtuA K43A |
| FN0900 | CACCATTATTACCAATAAAAACGGTC | |
| FN0901 | TTATTGGTAATAATGGTGCAGGTGCAACATCAATATTAAAATCG TTGGCAAC | |

| Primer | Sequence (5’ – 3’) | Description |
|---|---|---|
| FN0902 | GTTGCCAACGATTTTAATATTGATGTTGCACCTGCACCATTATT ACCAATAA | |
| FN0907 | ACATGCACCTGCATCCAACA | Amplification of pSeptuI for mutation PtuA D316A/E317A |
| FN0908 | TACAATTCCCTCGCCGTTTAGC | |
| FN0909 | AACGGCGAGGGAATTGTATTAATTGCTGCAGTGGACATGCACC TGCATCCAA | Amplification of pSeptuI for mutation PtuA H321A |
| FN0910 | TTGGATGCAGGTGCATGTCCACTGCAGCAATTAATACAATTCCC TCGCCGTT | |
| FN0923 | AACCGCCGGCTAGTTTGC | Amplification of pSeptuI for mutation PtuB K6A |
| FN0924 | GCGCATGTTTAATTTCCTGTTTGCG | |
| FN0925 | AGGAAATTAAACATGCGCGAAATAACTGCAGGGCAACCGCCGG CTAGTTTGC | |
| FN0926 | GCAAACTAGCCGGCGGTTGCCCTGCAGTTATTTCGCGCATGTT TAATTTCCT | |
| FN0919 | GCCAGGGATTTAGCTCCTG | Amplification of pSeptuI for mutation PtuB H75A |
| FN0920 | ACGGTATCGCGGTTTGTACC | |
| FN0921 | TACAAACCGCGATACCGTTAATGAAGCTGTCGAAGCCAGGGAT TTAGCTCCT | |
| FN0922 | AGGAGCTAAATCCCTGGCTTCGACAGCTTCATTAACGGTATCG CGGTTTGTA | |
